# Beyond the Classical Flake-to-Vesicle Mechanism: Non-standard Vesicles Arise from S-Shaped Intermediates

**DOI:** 10.64898/2026.11.24.768766

**Authors:** Zi-Kang Wang, Qian-Jun Xu, Jing-Wei Weng, Xu Xin

## Abstract

Vesicles are indispensable tools across numerous scientific and pharmaceutical fields, however, optimizing their properties necessitates a comprehensive understanding of their formation mechanism. The formation of lipid vesicles from bilayered lipid flakes has long been recognized as a fundamental step in vesicle preparation. The classical flake-to-vesicle mechanism, grounded in continuum models, describes this process as the transformation of a lipid flake into a regular spherical vesicle (RSV) via a cup-shaped intermediate. Nevertheless, this classical mechanism fails to account for emerging experimental observations of non-standard vesicles exhibiting diverse morphologies. Here, using extensive coarse-grained molecular dynamics simulations combined with Markov state model analysis, we systematically revisit this transition process at near-atomic resolution. In addition to the standard product, we observe a notable fraction of non-standard vesicles, including small-sized spherical vesicles (SSVs), ellipsoidal vesicles (ELVs), and flake-derived vesicle complexes (FVCs). These non-standard products arise from S-shaped intermediates that differ from the classical cup-shaped intermediate, and the degree of S-shapedness is closely correlated with whether non-standard vesicles are generated and, if so, which specific types. This correlation holds consistently across four lipid systems with distinct compositions and transbilayer distributions. On this basis, we propose a generalized mechanism that explicitly incorporates the S-shaped intermediates and the non-standard vesicles. This new mechanism opens new opportunities for the innovative development of engineered vesicles with tailored morphologies for diagnostics, drug delivery, and therapeutics.

## Introduction

Vesicles are closed structures formed by amphiphilic molecules, with an internal aqueous compartment. They can be assembled from naturally occurring lipids or from synthetic amphiphiles such as surfactants and block copolymers. Since the early 1960s^1,2^, vesicles have attracted considerable attention as model systems for biomembranes and as platforms for reconstituting membrane proteins. They have also found broad applications in biophysical studies, biochemical assays, and pharmaceutical development^3–5^. Clinically, vesicle-based formulations have been explored for ophthalmic disorders^4^, cancer^6,7^, and oral diseases^8^, as well as for vaccine delivery^9,10^, and gene therapy^11,12^. Despite these widespread applications, the mechanisms by which vesicles form from precursor structures remain incompletely understood, limiting rational design of their stability, reproducibility, and molecular encapsulation efficiency^13–15^.

The formation of lipid vesicles from bilayered lipid flakes (BLFs) has long been recognized as a fundamental step in various vesicle preparation techniques, including detergent depletion, ethanol injection, microfluidic hydrodynamic focusing, and stationary-phase interdiffusion.^3,16–20^ According to the classical flake-to-vesicle mechanism, the transition is primarily driven by the thermodynamic instability of the exposed hydrophobic edge of the BLF ^16,18,21,22^. To minimize the edge perimeter, the BLF bends, transiently adopts a cup-shaped structure, and ultimately seals to form a closed vesicle^16,18,22–24^.

Continuum models underpin the classical mechanism by treating the BLF as one or two smooth, structureless elastic sheets, characterizing the transition in terms of macroscopic quantities such as edge tension, bending modulus, and curvature radius^22–28^. Within this framework, the fate of a BLF is governed by the competition between edge energy, which favors the reduction of the exposed edge perimeter, and the bending energy, which penalizes membrane curvature. These continuum models have successfully rationalized the dependence of vesicle size on lipid composition, detergent type, and preparation rate^22,23^. Furthermore, the theoretically proposed cup-like intermediates have been experimentally corroborated by various technologies^29–33^, providing strong supports for the classical mechanism.

Despite these successes, accumulating experimental evidence challenges the universality of the classical mechanism, particularly regarding the emergence of non-standard vesicles. For instance, spontaneously formed ellipsoidal vesicles have been frequently observed^33–35^. While their stability can be accounted for by the area-difference-elasticity (ADE) model^36,37^, their formation mechanism remains ambiguous. Even more complicated morphologies, such as conjugated liposome assemblies^31,38^, have been reported, yet their formation mechanism remains largely elusive.

These discrepancies suggest that the classical flake-to-vesicle mechanism may be oversimplified, as the continuum models inherently neglect molecule-level details. Such details include intraleaflet lipid diffusion, lipid flip-flop (which relieves the interleaflet tension accumulated during bending), and the complex rearrangement of surrounding water and ions. Consequently, studies incorporating these neglected factors can provide a more realistic description of the process, holding the potential to bridge the gap between emerging experimental observations and the current flake-to-vesicle mechanism.

Molecular dynamics (MD) simulation is a valuable tool for providing simultaneous high-resolution spatial and temporal information on dynamic processes that are difficult to resolve with conventional experimental methods^39,40^. It can be regarded as a computational experiment in which dynamic processes and transient intermediates are directly “observed”^41^. Compared with the continuum models, MD simulation resolves chemical and biological processes at the level of individual lipid molecules and solvent, with atomistic/near-atomistic resolution^42,43^, and can therefore test whether the classical edge-energy/bending-energy balance correctly captures the driving force, identify the molecular intermediates that the continuum models averages out, and reveal how lipid tail order, headgroup hydration, lipid flip-flop, and solvent rearrangement shape the closure pathway. MD simulation therefore provides an opportunity to refine the classical flake-to-vesicle mechanism.

Using the planar bilayer as a simplified model of vesicles, Marrink *et al.* pioneered MD simulation of the self-assembly of randomly distributed lipids into planar bilayers by developing realistic molecular models for lipids^44^. By monitoring the self-assembly process at the atomistic/near-atomistic resolution, this work and its follow-up studies have pinpointed key intermediate states and provides mechanistic insights into the overall assembly pathway^44–47^. In our previous work, we combined MD simulation and Markov state model (MSM) analysis to derive free energy profiles and identify transition pathways, offering a succinct and physically meaningful description for the self-assembly process^47^. However, the planar bilayers obtained in all these simulation studies are constrained by periodic boundary conditions and a finite simulation box, and therefore lack key geometric and physical characteristics such as curvature, an encapsulated aqueous core, and asymmetric mechanical stress between the two leaflets.

To capture these missing features, the formation of intact spherical vesicles has also been investigated by MD simulations^15,17,48–55^. MD simulations may start from a random solution of lipid molecules at a very high concentration to accelerate self-assembly^17,48–52^, or may start directly from a BLF^15,53–55^. Nevertheless, a systematic investigation into the thermodynamics and kinetics of the flake-to-vesicle transition is still lacking. What is the free energy profile for this transition at the atomistic/near-atomistic scale? What mechanistic differences from the classical flake-to-vesicle mechanism does MD simulation reveal? How do lipid composition and transbilayer distribution influence the process? These questions remain unanswered.

In this work, we investigate the molecular mechanism of the flake-to-vesicle transition using MD simulations combined with MSM analysis. The Martini coarse-grained force field was employed to simulate the transition at near-atomistic resolution^56–59^, while MSM analysis was applied to derive thermodynamic and kinetic information from the trajectories^60–62^. This combination has been proved to be effective for deriving free energy profiles and identifying key intermediate states with appropriately selected collective variables^47^. Although the majority of MD trajectories support the classical mechanism by characterizing the flake-to-vesicle transition as a straightforward sequential process (flake → cup → vesicle)^16,18,22,23^ and yielding regular spherical vesicles (RSVs) as the product, a considerable number of S-shaped intermediates distinct from the cup-shaped ones are observed. Moreover, intermediates with a higher degree of S-shapedness give rise to non-standard products, including small-sized spherical vesicles (SSVs), ellipsoidal vesicles (ELVs), and flake-derived vesicle complexes (FVCs), and the degree of S-shapedness is closely correlated with the final product morphology. This correlation relationship is consistently observed across four lipid systems with distinct compositions and transbilayer distributions. Collectively, these results lead us to propose a generalized mechanism for the flake-to-vesicle transition that encompasses both the standard and non-standard products as well as the S-shaped intermediates.

## Methods

### MD simulation details

All systems were modeled using the Martini 2 coarse-grained force field^56,63–65^. The force field parameters were downloaded from the Martini website^65^. MD simulations were performed using GROMACS 2022.6, with trajectory extensions carried out using GROMACS 2025.3^66^. Detailed system preparation protocols^67^ are provided in the **Supplementary Materials**.

All production simulations were conducted in the isothermal–isobaric (NPT) ensemble at 310 K and 1 bar. Temperature was maintained using the V-rescale thermostat algorithm^68^ with a time constant of 1.0 ps. Pressure was controlled using the Parrinello–Rahman coupling algorithm^69^ with a time constant of 5.0 ps and a compressibility of 3e-4 bar^-1^. The equations of motion were integrated using the leap-frog algorithm with a time step of 20 fs. Electrostatic interactions were treated using the reaction-field method^70^ with a cutoff of 1.1 nm, a relative dielectric constant of 15, and an infinite dielectric constant for the reaction field. The Lennard–Jones interactions were truncated at 1.1 nm using the potential-shift-Verlet modifier. Periodic boundary conditions were applied to all three dimensions.

The total production simulation times for the four lipid systems are summarized in **Table 1**: 71.8 μs for the pure 1,2-dioleoyl-sn-glycero-3-phosphoethanolamine (DOPE) system, 22.5 μs the pure N-palmitoyl-D-erythro-sphingosyl-1-phosphocholine (DPSM) system, 80.0 μs for both the asymmetric plasma membrane (APM) and the symmetric plasma membrane (SPM) systems. Lipid coordinates were saved every 100 ps for subsequent analysis. Throughout this work, we report simulation times as the actual simulation time. Notably, the effective sampling time is four times the reported simulation time, owing to the accelerated dynamics of the Martini coarse-grained force field^42^ used in this study.

**Table 1.**
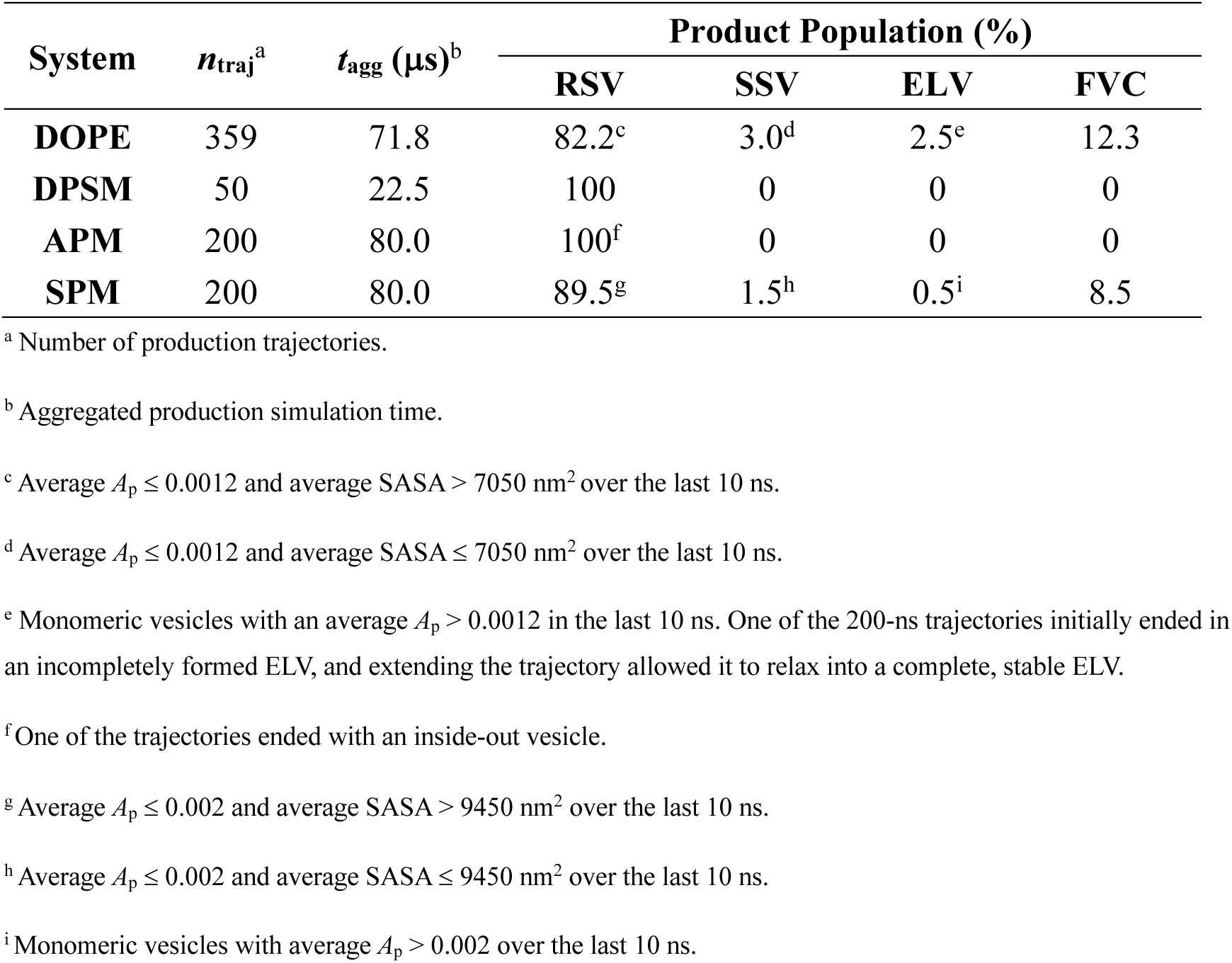
Number of trajectories yielding each product morphology for the systems with different lipid compositions and transbilayer distributions, including the pure 1,2-dioleoyl-sn-glycero-3-phosphoethanolamine (DOPE) system, the pure N-palmitoyl-D-erythro-sphingosyl-1-phosphocholine (DPSM) system, the asymmetric plasma membrane (APM) system, and the symmetric plasma membrane (SPM) system. Four product morphologies were identified: regular spherical vesicles (RSVs) and three non-standard types, such as small-sized spherical vesicles (SSVs), ellipsoidal vesicles (ELVs), and flake-derived vesicle complexes (FVCs).

Visualization of snapshots was performed using visual molecular dynamic (VMD)^71^.

### Calculation of S-shapedness index (SSI)

The S-shapedness index (SSI) was employed to quantify the S-shapedness of a lipid bilayer structure. The 2-D midsurface of a lipid bilayer structure (the surface located midway between the two leaflets) was represented as a triangular surface mesh, from which the mean curvature magnitude at each vertex and the boundary edge length were computed using our automated <u>a</u>nalysis f<u>r</u>amework for <u>c</u>urvature and bo<u>u</u>ndary geometry of membrane<u>s</u> (ARCUS) (see **Supplementary Materials** for details).

The concave and convex curvature magnitudes of the midsurface, denoted as *M*_concave_ and *M*_convex_, were computed by summing the weighted mean curvature magnitudes over the concave and convex regions, respectively:

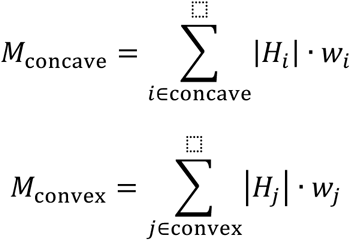

where |*H_i_*| and (*H*_j_(denote the mean curvature magnitudes at vertexes *i* and *j*, respectively, and *w_i_* and *w*_j_ denote the corresponding vertex weights of the vertexes, derived from the surface areas of the associated triangles. These values were calculated by the ARCUS approach. The summation indices run over all vertices assigned to the concave and convex regions, respectively.

The SSI is then defined as:

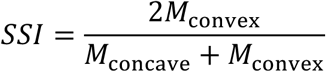

Concavity and convexity were assigned according to the orientation satisfying *M*_concave_ ≥ *M*_convex_. SSI = 0 indicates a typical cup-shaped structure, while SSI = 1 indicates a perfectly balanced S-shaped structure (*i.e.* half-concave, half-convex). In FVC-destined trajectories, once either daughter vesicle has formed, SSI cannot be calculated reliably, as the assignment of concave and convex regions becomes ambiguous.

The calculation of other features, including the asphericity parameter (*A*_p_), the solvent-accessible surface area (SASA) of lipids (**Figure S1**), the mean squared curvature, and the boundary edge length (**Figure S2**) are described in the **Supplementary Materials**.

### Markov state model (MSM) Analysis

MSM analysis was performed by the PyEMMA software package 2.5.6^72^. To build the MSM and compute the one-dimensional (1-D) free energy profile along the SASA of lipids, all the 295 RSV-destined flake-to-vesicle trajectories of the pure DOPE system were used. The first 10 ns of each trajectory were discarded as equilibrium, resulting in a total aggregate simulation time of 56.05 μs.

The structures from these trajectories were clustered into 100 microstates using the K-means algorithm^73^, with the lipid SASA as the sole clustering metric. This yielded a 100-state MSM. The lag time was determined to be 1.5 ns. The resulting MSM was rigorously validated by both the implied time scale test and the residence test (**Figure S3A,B**), and the mean first passage time (MFPT) estimated from the MSM was in good agreement with that directly obtained from the MD trajectories (95.07 ns; **Figure S3C,D**). In contrast, using *A*_p_ as the clustering metric failed to provide a valid MSM under the same validation criteria. The transition probability matrix obtained from the validated MSM was then used to compute the equilibrium distribution of the 100 microstates.

To construct the 1-D free energy profile, the lipid SASA values of all structures within each microstate were computed, and weighted by the inverse of the number of structures within the microstate and by the equilibrium distribution of the microstate. A weighted histogram (*N*(*SASA*)) was generated using 200 evenly spaced bins along the SASA coordinate. The free energy profile as a function of SASA, *F*(*SASA*), was then calculated as:

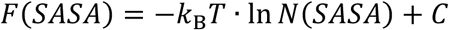

where *k*_B_ is the Boltzmann constant, *T* is the absolute temperature, and the constant *C* was chosen such that the free energy of the global minimum was zero.

Notably, the 1-D free energy profile was insensitive to the number of microstates in the MSM (**Figure S3E**).

## Results

### 82.2% of trajectories yield a regular spherical vesicle (RSV)

Using the MARTINI coarse-grained force field^56,63,64^, we simulated the flake-to-vesicle transition for a system composed of 6664 DOPE lipids (**Figure 1A**, **1B**). 359 independent 200-ns MD trajectories were collected (see **Supplementary Materials**), and most of them led to vesicle formation by the end of the simulations (**Table 1**). The asphericity parameter (*A*_p_) was employed to monitor the evolution of the trajectories (**Figure S1A**). A perfectly planar, infinitesimally thin disk corresponds to *A*_p_ = 0.25, whereas a perfect spherical vesicle corresponds to *A*_p_ = 0.^74,75^ As shown by the red curves in **Figure S1A**, the majority of the trajectories exhibit a distinct three-stage profile. Starting from initial BLF structures with *A*_p_ ≈ 0.24, *A*_p_ first undergoes a gentle decrease, then a steep decline, and finally plateaus below 0.0012, signifying the formation of spherical vesicles. Consistent with this, the solvent-accessible surface area (SASA) of lipids displays a similar three-stage profile, though with notably larger fluctuations. From an initial SASA of approximately 7850 nm^2^, most of the trajectories converge to a final state where SASA fluctuates around 7150 nm^2^ (red curves in **Figure S1B**). Visual inspection confirms that the end structures with *A*_p_ < 0.0012 and SASA > 7050 nm^2^ correspond to RSVs (**Figure 1C**). These end structures account for 82.2% of the total trajectories (**Table 1**) and are consistent with the final product morphology predicted by the classical flake-to-vesicle mechanism^15,16,23,49,50,52,53^.

**Figure 1.**
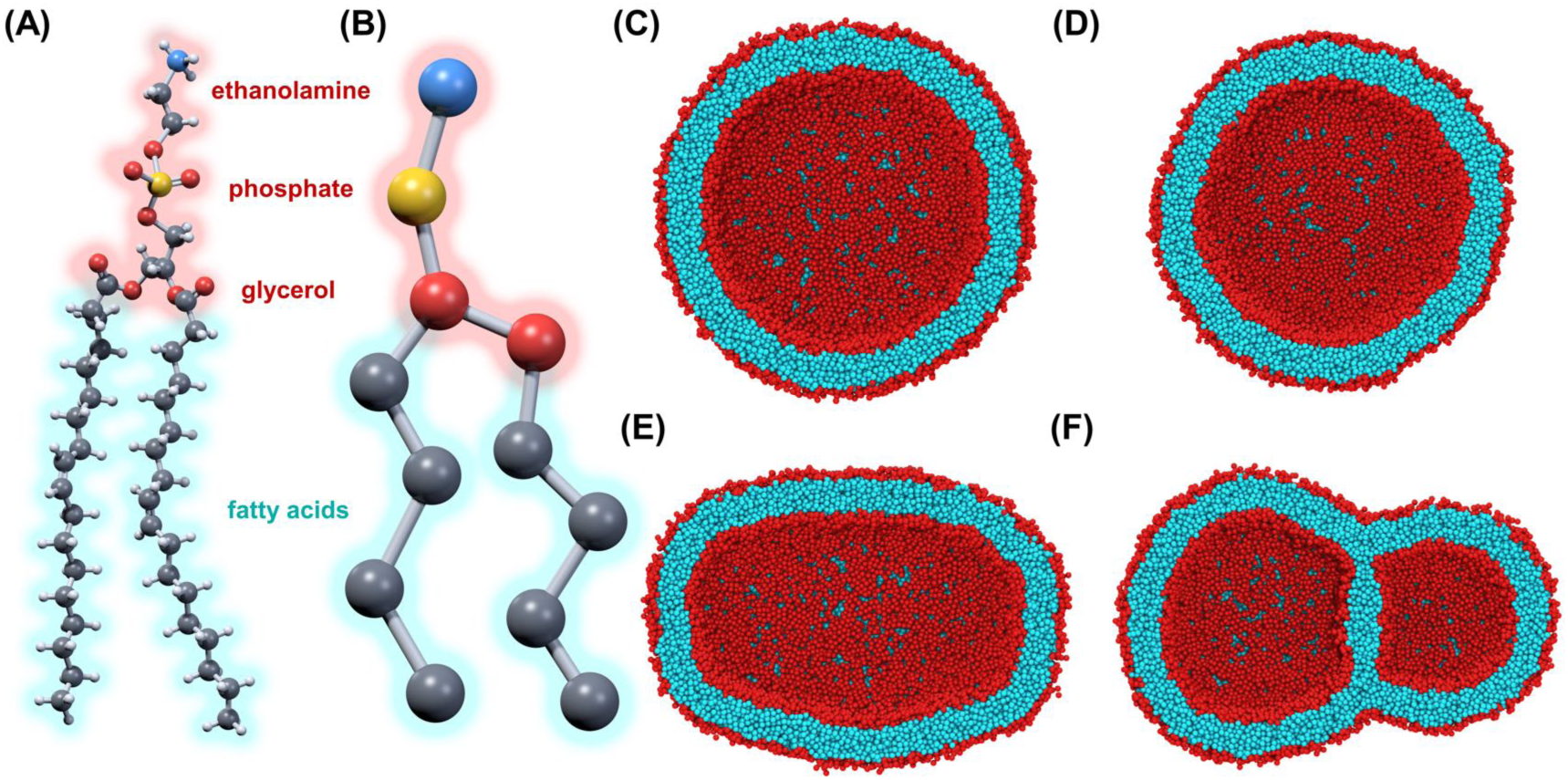
(A) Molecular structure of 1,2-dioleoyl-sn-glycero-3-phosphoethanolamine (DOPE). Atoms are colored by element (C: grey, H: white, N: blue, O: red, P: orange). (B) Coarse-grained representation of DOPE. (C-F) Representative snapshots of the different product morphologies observed in our simulations: (C) regular spherical vesicle (RSV), (D) small-sized spherical vesicle (SSV), (E) ellipsoidal vesicle (ELV), and (F) flake-derived vesicle complex (FVC).

To elucidate the transition process in more detail, we developed an automated <u>a</u>nalysis framework for <u>c</u>urvature and bo<u>u</u>ndary geometry of membrane<u>s</u> (ARCUS). This protocol approximates the lipid bilayer as a two-dimensional (2-D) midsurface, consistent with Helfrich theory^26^, and allows computing both the mean squared curvature and the boundary edge length (**Figure S2**; see **Supplementary Materials** for details). The mean squared curvature exhibits a three-stage profile (**Figure S2A**) that closely parallels the three-stage patterns of *A*_p_ and SASA (**Figure S1**), although it changes in the opposite direction: curvature increases as the vesicle closes, whereas *A*_p_ and SASA decrease. The boundary edge length also follows a three-stage profile, but its trend is opposite to that of the mean squared curvature: it decreases from an initially large value to zero at the final stage (**Figure S2B**). This is consistent with the notion of the continuum models that boundary edge energy contributes to driving the transition^18,22,24^.

Notably, a fraction of trajectories (17.8%, **Table 1**) end with a non-zero *A*_p_ between 0.0012 and 0.14 (**Figure S1A**) or with a SASA value below 7050 nm^2^ (**Figure S1B**), indicating the formation of non-standard vesicles such as SSVs, ELVs and FVCs (**Figure 1D-F**), which clearly deviate from the classical mechanism. We begin our analysis with RSVs and then turn to these non-standard vesicles in the following sections.

### RSV formation: similarities to and deviations from the classical flake-to-vesicle mechanism

For a rigorous yet succinct description of the flake-to-vesicle process, we used the RSV-destined trajectories (82.2% of all trajectories) to construct a validated MSM with 100 states (**Figure S3**), and derived a one-dimensional (1-D) free energy profile using the lipid SASA as the collective variable (see **Methods**). The 1-D free energy profile exhibits two free energy minima separated by a barrier (**Figure 2A**), characteristic of a two-state transition. The stable flake state corresponds to the minimum at SASA = 7837 nm^2^, while the RSV state corresponds to the other at SASA = 7153 nm^2^ (**Figure 2A, S1B**). The RSV state lay 29.5 kJ/mol lower in free energy than the stable flake state, indicating that vesicle formation is thermodynamically favorable (**Figure 2A**).

**Figure 2.**
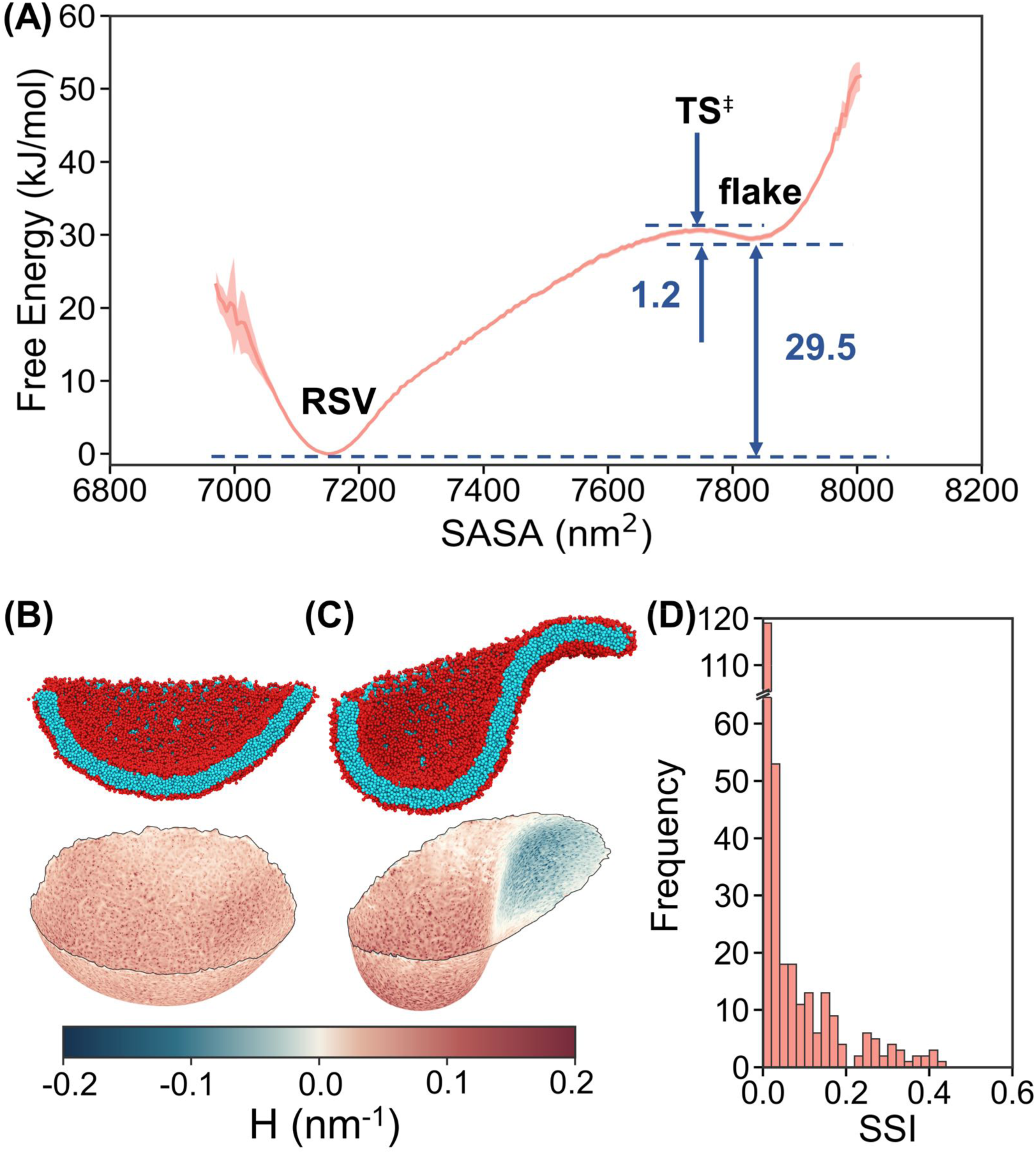
(A) One-dimensional (1-D) free energy profile as a function of the solvent-accessible surface area (SASA) of lipids for the flake-to-vesicle transition of the pure DOPE system. Only RSV-destined trajectories were used in the calculation. Shaded areas represent the standard error estimated via bootstrapping. (B) Cross-sectional snapshot and corresponding 2-D midsurface of a cup-shaped intermediate with SSI = 0. Hydrophobic and hydrophilic segments of DOPE lipids are colored cyan and red in the cross-sectional snapshot, respectively. The midsurface is colored by mean curvature *H*. (C) Cross-sectional snapshot and corresponding 2-D midsurface of an S-shaped intermediate with SSI = 0.42. (D) Frequency distribution of the SSI for the key-progress-position (KPP) intermediate of each trajectory that successfully reached an RSV.

The 1-D free energy profile reveals that the stable flake state is separated from the RSV state by a free energy barrier of 1.2 kJ/mol (**Figure 2A**). The transition state, located at 17% along the overall transition progress, corresponds to a SASA of 7748 nm^2^. The presence of this barrier is consistent with the three-stage profiles of the monitored variables, such as *A*_p_ and SASA (**Figure S1, S2**). Initially, the system evolves slowly along a shallow free energy slope, corresponding to the gentle stage. After surmounting the barrier, it rapidly relaxes toward the RSV state along the downhill free energy gradient, as indicated by the steep stage. It finally reaches the RSV state, where the monitored variables remain nearly constant, corresponding to the plateau stage. Notably, the presence of this barrier is broadly consistent with analytical estimates from the classical phenomenological model, which predicts an energy barrier associated with a cup-shaped intermediate^23^.

Despite the above consistency, our MD simulations revealed unexpected deviations from the classical mechanism. While the classical mechanism proposes a predominant cup-shaped intermediate^16,23^, our simulations identified a considerable population of S-shaped intermediates, characterized by the coexistence of concave and convex regions within a single structure. To quantify the degree of S-shapedness of these structures, we defined the S-shaped index (SSI) using the mean curvatures calculated *via* the ARCUS approach (see **Methods**). In this metric, SSI = 0 corresponds to a typical cup-shaped morphology (**Figure 2B**), whereas SSI = 1 denotes a perfectly balanced S-shaped structure (*i.e.*, half-concave, half-convex). SSI values between 0 and 1 therefore represent unbalanced S-shaped structures with a greater proportion of concave regions (**Figure 2C**).

For further analysis, the intermediate structures at 17% of the overall transition progress are used as representative intermediates and are referred to as key-progress-position (KPP) intermediates, as they lie near the transition-state region of the RSV-destined trajectories (**Figure 2A**). The SSI distribution of these KPP structures underscores the structural heterogeneity of intermediates (**Figure 2D**). Although 74.2% of these structures adopted morphologies closely resembling the cup-shaped geometry (with SSI values < 0.1), the remaining 25.8% exhibited more distinct S-shape morphologies with larger SSI values, some reaching as high as 0.42 (**Figure 2D**). Such high SSI values indicate the presence of convex regions in the intermediates, and reflect the intrinsically high flexibility of lipid bilayers, a feature neglected by the classical mechanism^23^. Although these intermediates suggest additional transition pathways and can modestly accelerate the transition, their relatively low proportion limits their overall impact on the RSV-forming process.

### Three types of non-standard vesicles generated from a bilayered lipid flake

Although BLFs transform into RSVs in the majority of MD trajectories, the remaining 17.8% of trajectories yield three distinct types of non-standard vesicles: SSVs, ELVs, and FVCs (**Table 1**; **Figure 1D-F**). While these non-standard vesicles have higher free energy than RSVs, they are kinetically stable, appearing as local minima on the free energy landscape (**Figure S4**).

SSVs, which account for 3.0% of all trajectories (**Table 1**), are spherical (*A*_p_ < 0.0012) but consistently smaller than RSVs (**Figure 1D**). This is evidenced by their smaller inner and outer leaflet radii (**Figure 3A**) and lower lipid SASA (**Figure S1B**). This size reduction arises because SSVs encapsulate only 95% of the solvent content (water and ions) of RSVs (**Figure 3B**), while maintaining a lipid ratio between the outer and inner leaflets similar to that of RSVs (**Figure 3C**). Given limited water permeation (**Figure S5**) and the absence of lipid flip-flop across the bilayer, the reduced internal volume is trapped and the smaller spherical shape is stabilized even when the simulation time is extended to 1 μs (blue curves in **Figure S6A, S6B**).

**Figure 3.**
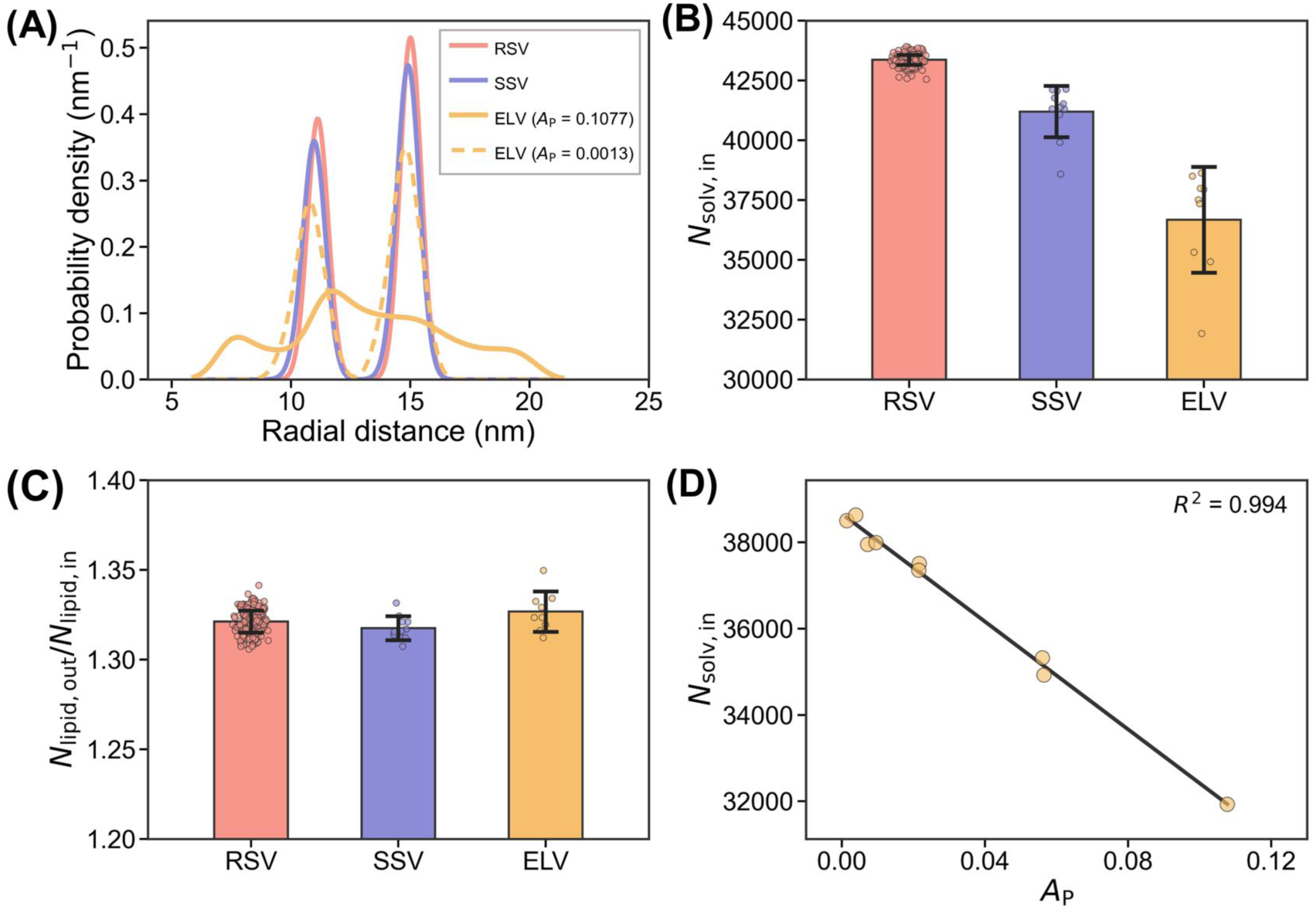
(A) Radial distributions of phosphate beads of lipids in the inner and outer leaflets of an RSV, an SSV, and two ELVs with *A*_p_ of 0.1077 and 0.0013, respectively. (B) Average number of internal solvent beads (*N*_solv,in_) in RSVs, SSVs, and ELVs. (C) Average lipid ratio between the outer and inner leaflets in RSVs, SSVs and ELVs. (D) Correlation between the number of internal solvent beads and *A*_p_ of ELVs. All data shown are for the pure DOPE system.

ELVs account for 2.5% of all trajectories (**Table 1**) and are characterized by aspherical shapes (**Figure 1E**), with *A*_p_ values ranging from 0.0013 to 0.1077, corresponding to shapes from quasi-spherical to highly elongated (**Figure S7**). Despite this morphological diversity, ELVs remain stable on a microsecond timescale (orange curves in **Figure S6A,B**). Like SSVs, ELVs maintain an outer-to-inner leaflet lipid ratio comparable to that of RSVs (**Figure 3C**), yet they enclose significantly fewer solvent beads (**Figure 3B, S8**). More importantly, we observed a strong correlation, across ELVs, between the number of internal solvent beads and the *A*_p_ value (**Figure 3D**), suggesting that the internal solvent content plays a key role in determining their shape.

In support of this, separate simulations demonstrate the causal role of internal solvent content in determining the shape of SSVs and ELVs. Removing internal solvent beads from an RSV induces its transformation into either an SSV (blue curves in **Figure S9**) or an ELV (orange curves in **Figure S9**), depending on the number of beads removed. An SSV is obtained when approximately 2200 internal solvent beads are removed, while an ELV is obtained when approximately 6700 are removed.

This morphological change is consistent with the ADE model, which posits that the shape of a vesicle is determined by minimizing the bending energy of its bilayer under the constraints on the membrane surface area and the internal solvent volume *V*^36,37^. The shape is then governed by two dimensionless parameters: the reduced volume *v* = 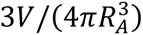, where *R_A_* = (*A*/4*π*)^1/2^ is the area-equivalent radius, and the effective spontaneous curvature *c̅*_0_, which depends on the area difference between the two leaflets. Because SSVs and ELVs share the same total number of lipids and a similar lipid ratio between the outer and inner leaflets (**Figure 3C**), they have similar surface areas and interleaflet area differences, and hence similar *R*_A_ and *c̅*_0_, but differ in *V*. When the vesicle encloses a volume close to the maximum that can be achieved by the given membrane surface area, *v* is close to 1, and the vesicle largely maintains a spherical shape, as in the case of SSVs. As *v* decreases below 1, the vesicle deviates from sphericity and becomes increasingly elongated, as in the case of ELVs (**Figure 3D**).

An FVC differs fundamentally from the monomeric vesicles described above, as it consists of two small daughter vesicles connected by a three-way junction, with a mutual bilayer separating the internal contents of the two daughter vesicles (**Figure 1F**). The relative sizes of the daughter vesicles range from markedly different to nearly comparable, although the former is less common (corresponding to FVC-destined trajectories with relatively low *A*_p_ in **Figure S1A**). The FVCs show considerable structural similarities to the conventional hemifused state that arises as an unstable intermediate during the fusion of two isolated vesicles: the daughter vesicles correspond to the hemifused vesicles, the mutual bilayer represents the hemifusion diaphragm (HD), and the three-way junction aligns with the HD rim^41^. The conjugated liposome assemblies reported by Kotoucek *et al.* share the same structural features as the FVCs and the hemifused state^38^. Interestingly, unlike the conventional hemifused states observed in previous MD simulations, which exhibit a maximal lifetime on the microsecond timescale^76–78^, the FVCs remain remarkably stable even when the simulation time is extended to 25 μs (**Figure S6C, S6D**), which corresponds to an effective 100 μs if a 4-fold timescale normalization is applied^77^.

### Correlation between the final product morphology and the S-shapedness of intermediates

Although non-standard vesicles are observed as final products, the underlying formation mechanism remains unclear: how do these vesicles arise, and why does a BLF transform into a non-standard vesicle instead of an RSV? To address these questions, we projected all trajectories onto a 2-D space defined by lipid SASA and SSI (**Figure 4A**): the decrease in lipid SASA tracks the progressive closure of the initial BLF into a vesicle, while SSI reflects the S-shapedness of individual snapshots.

**Figure 4.**
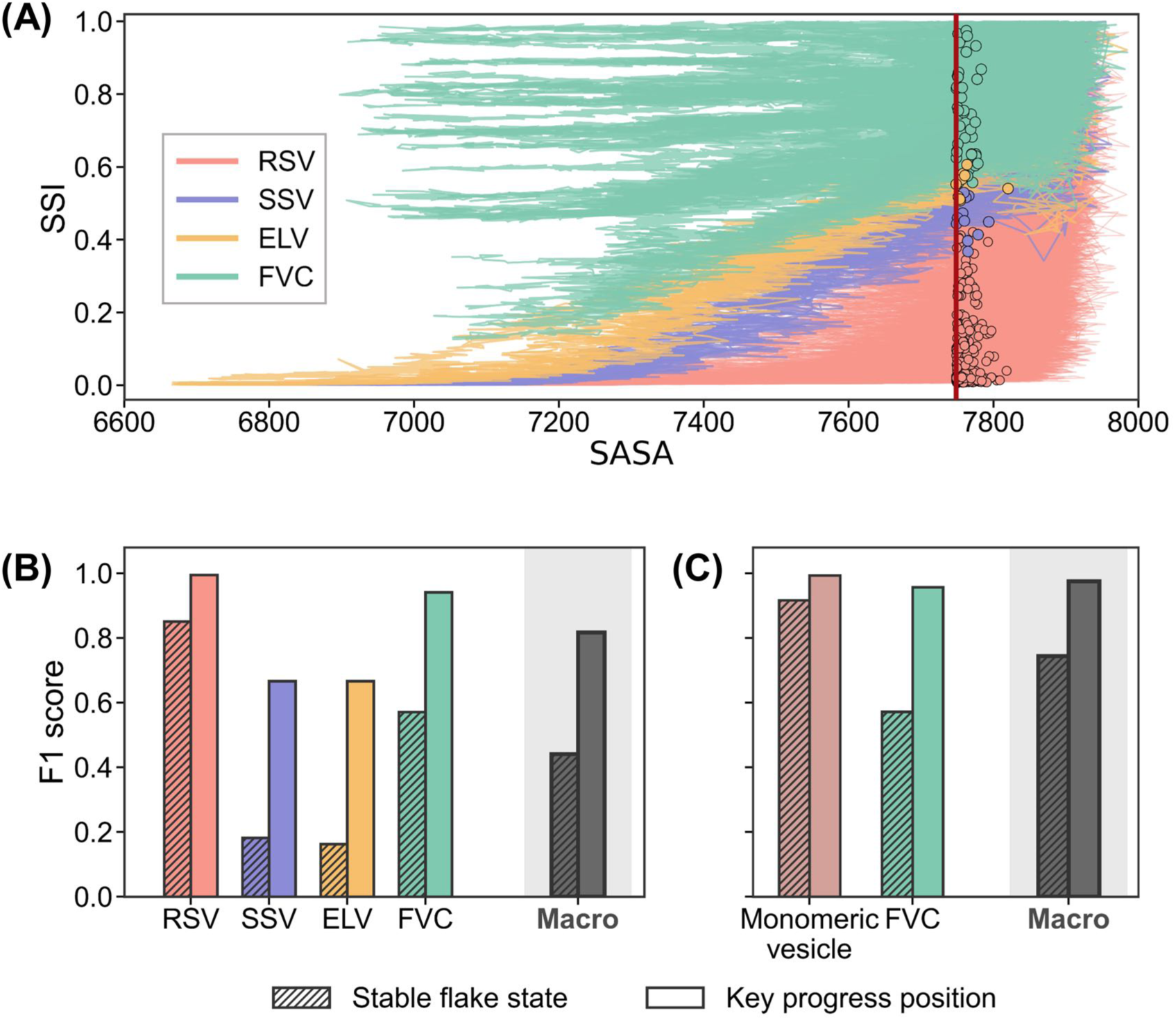
(A) Projection of all 200-ns trajectories of the pure DOPE system onto the 2-D space defined by lipid SASA and SSI. Trajectories are colored by their final product morphology. The snapshot immediately preceding the successful crossing of the key progress position in each trajectory is used to represent the KPP intermediate and is highlighted as a circle. FVC-destined trajectories are truncated after the formation of the three-way junction that greatly complicates the definition of SSI. (B) Per-class and macro-averaged F1-scores for the classification of the four final product morphologies using SSI_KPP_ or SSI_SFS_ as the sole feature (KPP = key progress position, SFS = stable flake state). (C) Per-class and macro-averaged F1-scores for the binary classification of monomeric vesicles versus FVCs using SSI_KPP_ or SSI_SFS_ as the single feature.

For RSV-destined trajectories, although the stable flake state samples a broad range of S-shaped conformations, with the SSI of the stable flake state (SSI_SFS_) fluctuating between 0 and 1 (red lines in **Figure 4A**), the SSI of intermediates at the key progress position (SSI_KPP_) is confined to a narrower range of 0–0.42 (red circles in **Figure 4A**; see also **Figure 2D** for the distribution). This narrowing suggests that successful evolution towards RSVs is associated with a specific range of SSI_KPP_. Similarly, the KPP intermediates of SSVs, ELVs, and FVCs occupy largely distinct SSI_KPP_ ranges, each being characteristic of the corresponding product (blue, orange, and cyan circles in **Figure 4A**, respectively). Thus, the MD trajectories lead to different products by passing through distinct SSI_KPP_ ranges.

These observations raise the question of whether SSI_KPP_ alone is sufficient to distinguish the final product morphologies. To test this, we used SSI_KPP_ as the sole feature for classifying the four product morphologies (see **Supplementary Materials**). With optimized thresholds, the per-class F1-scores for RSVs, SSVs, ELVs and FVCs reached 0.995, 0.667, 0.667 and 0.941, respectively, yielding a macro-averaged F1-score of 0.817 (**Figure 4B**). When the task was reduced to binary classification, *i.e.* distinguishing monomeric vesicles from FVCs, both per-class and macro-averaged F1-scores further improved to above 0.975 (**Figure 4C**). These results demonstrate that SSI_KPP_ alone can discriminate the final product morphologies with high accuracy, particularly for RSVs and FVCs.

For comparison, we evaluated the discriminatory power of SSI_SFS_ using the same classification framework. The macro-averaged F1-scores of SSI_SFS_ dropped to 0.441 and 0.744 for the four-class and two-class tasks, respectively (**Figure 4B**, **4C**). The lower performance of SSI_SFS_ suggests that the stable flake state carries less discriminatory information about the final product morphology, possibly due to its thermal fluctuations.

A representative RSV-destined trajectory illustrates the formation of an RSV from a non-classical S-shaped intermediate. As the BLF curled, a KPP intermediate with an SSI of 0.070 formed, whose shape slightly deviated from the cup shape (**Figure 5A**). Driven continuously by edge tension, the curling process involved two concurrent events: the concave region continuously increased its curvature, progressively evolving toward a spherical morphology, while the initially small convex region underwent curvature inversion, becoming concave and further increasing its curvature (**Figure 5B**, SSI = 0.008). Just prior to edge closure, the curvatures of almost all regions had converged to those of the final RSV (**Figure 5C**). The intermediate ultimately closed into a spherical vesicle containing a large number of solvent beads (**Figure 1C**).

**Figure 5.**
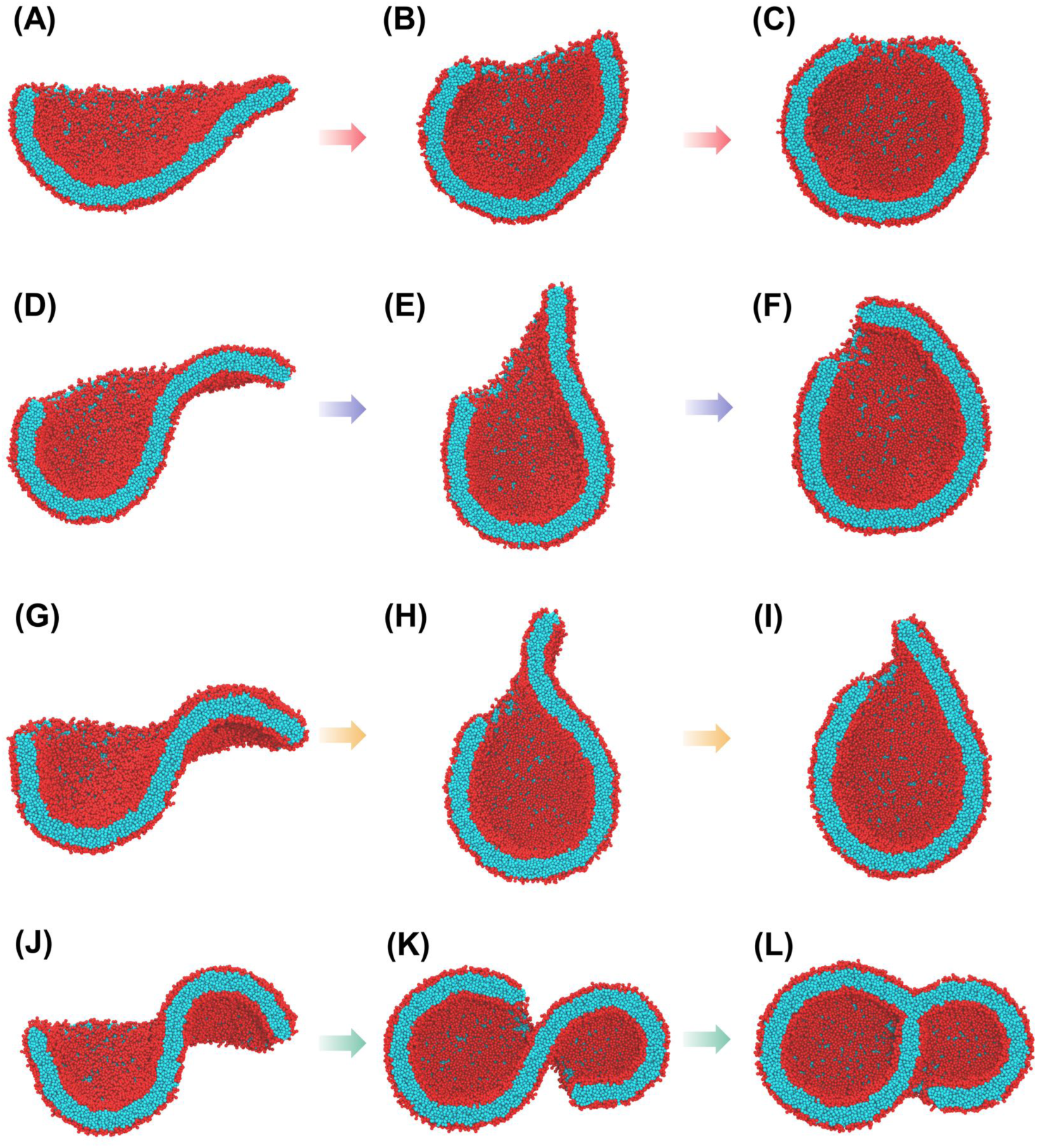
Cross-sectional snapshots of representative intermediates along the formation pathways of (A-C) RSV, (D-F) SSV, (G-I) ELV, and (J-L) FVC composed of pure DOPE. For each pathway, three key structures are shown: the key-progress-point (KPP) intermediate (A, D, G, J), the pre-closure intermediate (C, F, I, L), and the instantaneous structure between the two (B, E, H, K). Hydrophobic and hydrophilic segments of lipids are colored in red and cyan, respectively.

The formation of an SSV differs in one key aspect: the S-shaped intermediate retained a more pronounced convex region, as reflected by a larger SSI of 0.460 (**Figure 5D**). This convex region evolved towards concavity (**Figure 5E**; SSI = 0.046), but was kinetically trapped before edge closure, preventing it from fully equilibrating. The pre-closure intermediate was slightly aspherical and enclosed fewer solvent beads than the corresponding RSV intermediate (**Figure 5F**; SSI = 0.006). As discussed above, the reduced internal solvent content determines the smaller size of the final SSV morphology after equilibration (**Figure 1D**, **3B, S9**).

During the formation of an ELV, the KPP intermediate exhibited a more pronounced S-shaped morphology than that observed for SSVs (**Figure 5G**; SSI = 0.553). As the transition proceeded (**Figure 5H**; SSI = 0.080), this larger convexity resulted in a greater deviation of the local curvature from that of an RSV. Crucially, although the convex-to-concave inversion was largely completed in the pre-closure intermediate (**Figure 5I**; SSI = 0.015), the system remained kinetically trapped, as in the SSV case, and its curvature deviated from that of an RSV more than in the SSV case. At the pre-closure stage, the intermediate was more elongated than that of an SSV and therefore enclosed less solvent. After closure, this reduced solvent content was trapped and stabilized the ellipsoidal morphology (**Figure 1E**) rather than a spherical one (**Figure 3B**).

FVCs represent the tipping point, where a quantitative increase in SSI_KPP_ gives rise to a qualitative shift in the formation pathway. As the convex region enlarged further, with an even higher SSI_KPP_ of 0.821 (**Figure 5J**), the intermediate no longer followed the convex-to-concave pathway. Instead, the convex and concave regions curled independently (**Figure 5K**) and their boundary edges fused with the adjacent bilayer, driven by edge tension. This ultimately yielded two small daughter vesicles separated by the mutual bilayer (**Figure 1F**). The relative size of the two daughter vesicles was closely related to SSI_KPP_: an intermediate with a concave region larger than its convex region (*i.e.*, a relatively smaller SSI_KPP_ among FVC-destined trajectories) evolved into an FVC with a larger daughter vesicle conjoined with a smaller one, whereas an intermediate with comparable concave and convex regions (a relatively larger SSI_KPP_) produced two daughter vesicles of similar size. Further inspection showed that the larger daughter vesicle closed earlier than the smaller one in most of the trajectories (**Figure 5L**). Edge tension drives the closing of the two daughter vesicles, and the resulting FVC is kinetically stabilized against reverting to the S-shaped intermediate.

Collectively, these snapshots reveal four distinct transition pathways from S-shaped intermediates with varying SSI_KPP_ values to their corresponding final products (**Figure 5**). This is consistent with the distinct SSI_KPP_ ranges identified for each product morphology (Figure 4A), reinforcing the correlation between the final product morphology and SSI_KPP_.

### Conservation of the morphology–S-shapedness correlation in three other lipid systems

DOPE serves as a canonical example of a lipid with a strong negative monolayer spontaneous curvature, which arises from its relatively small headgroup and bulky hydrophobic tails (**Figure 1A,B**). Although the preceding results for the DOPE bilayer system have established the existence of S-shaped intermediates and non-standard vesicles, as well as the correlation between them, a critical question remains: are these phenomena specific to the DOPE system, or do they extend to bilayer systems with different compositions or transbilayer distributions? To test the generality of our findings, we examined three additional lipid systems.

We first selected a system composed purely of DPSM, a lipid with a positive monolayer spontaneous curvature due to its relatively large headgroup and slender tails (**Figure 6A**, inset). All 50 flake-to-vesicle trajectories of the pure DPSM system (see **Methods**) terminated in RSVs, with no non-standard vesicles observed (**Table 1**; **Figure 6A, S10**). Although the flake structures transiently sampled S-shaped conformations, with their SSI values fluctuating between 0 and 1 (**Figure 6A**), all SSI_KPP_ values converged to approximately zero (red circles, **Figure 6A**), with the key progress position still defined at 17% of the overall progress, as in the pure DOPE system. This indicates that the correlation between SSI_KPP_ and the final product morphology is also maintained in the DPSM system, in which intermediates with small SSI_KPP_ values evolve into RSVs. The absence of S-shaped intermediates with large SSI_KPP_ likely arises from the substantially higher bending modulus of the DPSM bilayer^79^, which imposes a high energy penalty on complex morphologies such as the S-shape.

**Figure 6.**
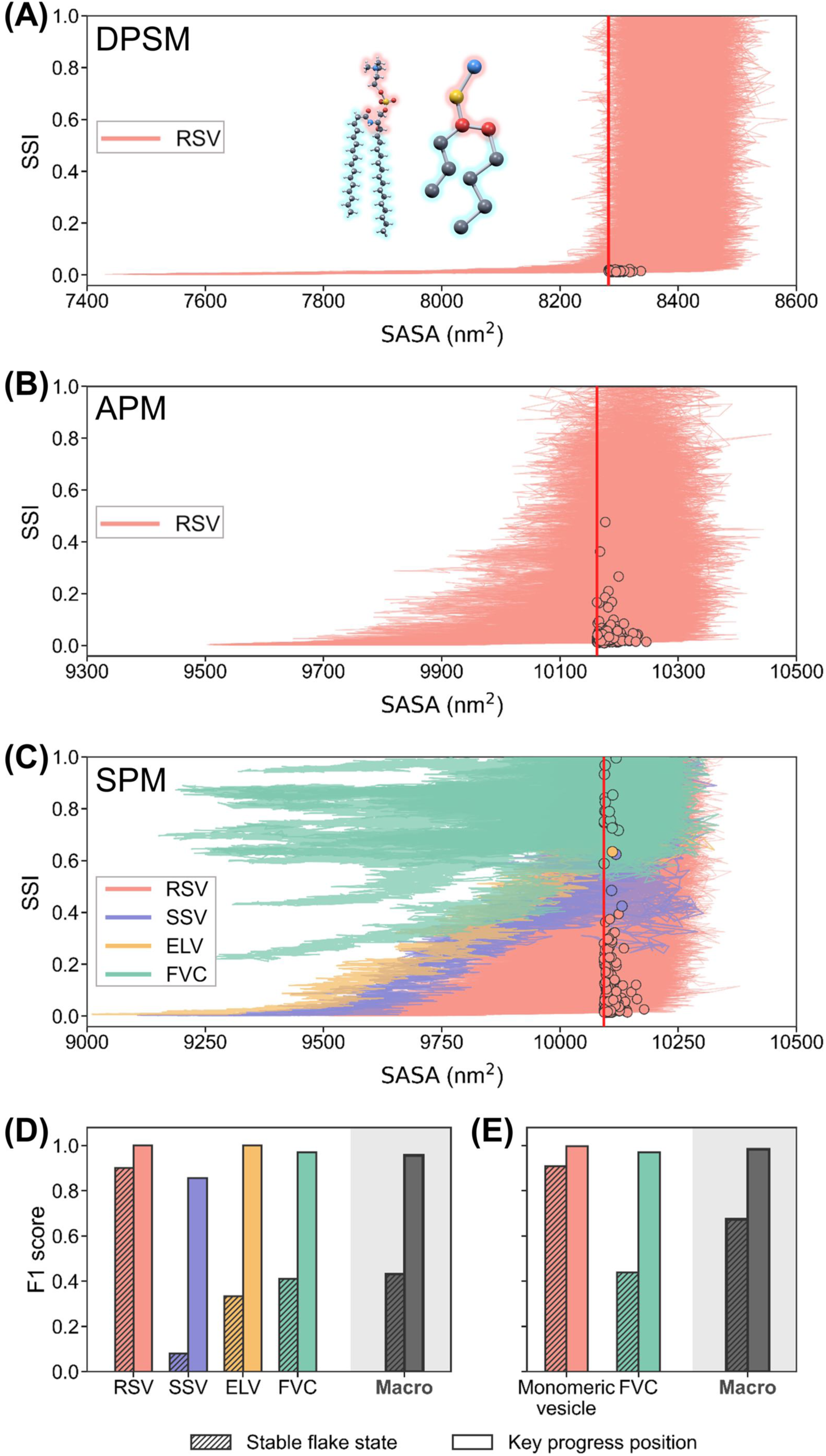
Projection of the flake-to-vesicle trajectories onto the 2-D space defined by lipid SASA and SSI for (A) the pure N-palmitoyl-D-erythro-sphingosyl-1-phosphocholine (DPSM) system, (B) the asymmetric plasma membrane (APM) system with asymmetrically distributed lipids across two leaflets, and (C) the symmetric plasma membrane (SPM) system with symmetrically distributed lipids. Data points are colored by the final product morphology: red for RSVs, blue for SSVs, orange for ELVs, and cyan for FVCs. The vertical red line indicates the key progress position. The snapshot immediately preceding the successful crossing of the key progress position in each trajectory is used to represent the KPP intermediate and is highlighted as a circle. (D) Per-class and macro-averaged F1-scores for the classification of the four final product morphologies of the SPM system using SSI_KPP_ as the sole feature. The ELV class contains only a single trajectory, and its F1-score should therefore be interpreted with caution. (E) Per-class and macro-averaged F1-scores for the binary classification of monomeric vesicles versus FVCs using SSI_KPP_ as the solo feature in the SPM system.

Cell-membrane-coated biomimetic vesicles have recently emerged as nanocarriers that provide unique physiological functions for drug delivery^80^. To investigate the formation mechanism of such biomimetic vesicles, we employed a plasma membrane model composed of 63 distinct lipid species with 14 headgroup types and 11 tail types^57^. In the initial BLF, lipids are symmetrically distributed across the two leaflets, and this system is referred to as the symmetric plasma membrane (SPM) system. For comparison, we also prepared the asymmetric plasma membrane (APM) system, in which lipids are asymmetrically distributed across the two leaflets, mimicking the native plasma membrane^57^ (see **Supplementary Materials**). The APM system was used to assess the role of transbilayer distribution in the vesiculation process.

All trajectories of the APM system resulted in RSVs (**Table 1**; **Figure 6B, S11**). Because the asymmetric transbilayer distribution generates a spontaneous curvature, cup-shaped intermediates are favored, whereas highly S-shaped structures are suppressed in the APM system. Under the assumption that the APM system has the same key progress position as that of the pure DOPE system (red vertical line in **Figure 6B**), we calculated SSI_KPP_ values (red circles in **Figure 6B**), which are small and whose range is generally similar to that of RSV-destined DOPE trajectories (red circles in **Figure 4A**). These results corroborate the correlation between the final product morphology and SSI_KPP_ in a highly complex and asymmetric system.

Despite having a lipid composition similar to that of the APM system, the SPM system exhibited markedly higher morphological diversity, yielding all four product morphologies (RSVs, SSVs, ELVs, and FVCs; **Table 1**; **Figure S12**). Since the two systems share a similar lipid composition but differ in transbilayer distribution, this contrast indicates that transbilayer lipid distribution plays a critical role in determining the final product morphology. Compared with the pure DOPE system, RSVs remain the most abundant product (89.5%) in the SPM system, whereas all non-standard vesicles appear at lower frequencies (**Table 1**). The greater similarity in product morphological diversity between the SPM and pure DOPE systems, rather than between the SPM and DPSM systems, is likely due to the considerable proportion of lipids with negative monolayer spontaneous curvature in the SPM system: 16% with small phosphatidylethanolamine (PE) headgroups and approximately 30% with bulky tails containing multiple unsaturated bonds^57^.

To assess the discriminatory power of SSI_KPP_ for the SPM system, we adopted the same assumption regarding the key progress position (red vertical line in **Figure 6C**) and calculated the corresponding F1-scores. For the four-class classification task, the per-class F1-scores remain high: 1.0 for RSVs, 0.857 for SSVs, 1.0 for the sole ELV, and 0.970 for FVCs (**Figure 6D**). For the binary classification task distinguishing monomeric vesicles from FVCs, both the per-class and macro-averaged F1-scores exceed 0.970 (**Figure 6E**). These high F1-scores demonstrate that the compositional complexity of the SPM system does not diminish the discriminatory power of SSI_KPP_. Together with the DPSM and APM systems, these results establish the conservation of the morphology–S-shapedness correlation across three other lipid systems.

## Discussion

Vesicles have emerged as a critical class of tools with broad applications across numerous scientific disciplines and the pharmaceutical industry^3–5^. A deep understanding of the formation mechanism of vesicles is crucial for improving their stability, reproducibility, and molecular encapsulation efficiency^23,24^.

The formation of lipid vesicles from BLFs has long been recognized as a fundamental step in vesicle preparation^3,16–20^. According to the classical mechanism, this transition proceeds from a BLF through a cup-shaped intermediate, ultimately yielding a RSV. Although this mechanism is supported by extensive experimental evidence^29–33^, it cannot fully account for the experimentally observed non-standard vesicles, such as ellipsoidal vesicles^31,34,35,38^ and conjugated liposome assemblies^38^.

In this study, we employ coarse-grained MD simulation combined with MSM analysis^60,61^ to characterize the flake-to-vesicle transition at near-atomistic resolution. Although the MARTINI coarse-grained force field^56,63,64,81^ inherently offers only semi-quantitative accuracy, our simulations of the pure DOPE system successfully reproduce RSVs as the major product (**Figure 1C** and **Table 1**) and elucidate the flake-cup-RSV sequence as the dominant pathway, providing robust support for the classical mechanism. Crucially, however, a notable subset of trajectories captures non-canonical events, culminating in diverse non-standard vesicles such as SSVs, ELVs, and FVCs (**Table 1**; **Figure 1D-F**).

Like RSVs, both SSVs and ELVs constitute monomeric vesicles. While SSVs retain a spherical conformation despite their reduced size (**Figure 1D**), ELVs exhibit aspherical conformations, spanning from quasi-spherical to highly elongated geometries (**Figure 1E, S7**). These morphological deviations from the canonical RSV shape stem primarily from the internal solvent content at the instant of vesicle closure. Specifically, a higher internal solvent content promotes spherical geometries, whereas a lower content yields aspherical morphologies^36,37^ (**Figure 3B**, **3D, S9**). Although spontaneously formed ELVs have been frequently observed in experiments^33–35^, their formation mechanism has remained ambiguous. Our work resolves this ambiguity by demonstrating that stable ELVs can be generated directly from lipid flakes, thereby providing a mechanistic explanation for their experimental observations.

In contrast to monomeric vesicles (RSVs, SSVs, and ELVs), the FVCs observed in our simulations are structurally distinct. Each FVC comprises two daughter vesicles connected by a three-way junction, with their internal contents separated by a mutual bilayer (**Figure 1F**). This architecture exhibits considerable structural similarities to both the conventional hemifused state arising from the fusion of two isolated vesicles^41^ and the conjugated liposome assemblies reported by Kotoucek et al^38^. However, both the conventional hemifused state and the conjugated liposome assemblies are considered short-lived intermediates^38,41^. Specifically, the conventional hemifused state has a maximal lifetime on the microsecond timescale in MD simulations^76–78^, whereas our FVCs exhibit stability on the submicrosecond timescale (**Figure S6C, S6D**). This discrepancy can be reconciled by the recent discovery of hemifusomes, a novel class of vesicular organelle complexes that share structural features with the conventional hemifused state^82^ but possess exceptionally long lifetimes. Their existence suggests that hemifused-state-like structures can be highly stable. Further experiments are required to determine whether BLFs can, under appropriate conditions, produce FVCs with hemifusomes-like longevity.

The identification of S-shaped intermediates represents another key finding of this work, arising from our efforts to trace the origins of the final products. In the pure DOPE system, four distinct transition pathways were observed, in which different final product morphologies arise from intermediates with various degrees of S-shapedness (**Figure 5**). Furthermore, the SSI of intermediates shows strong discriminative power for the final product morphology (**Figure 4B**, **4C**). These results demonstrate the close correlation between the final product morphology and the S-shapedness of intermediates. Remarkably, this morphology-S-shapedness correlation is retained in other three lipid systems with various lipid compositions and transbilayer distributions. The pure DPSM and APM systems^57^ have intermediates of a low degree of S-shapedness and form only RSVs (**Figure 6A**, **6B**). The SPM system generates all four types of vesicles, and the degree of S-shapedness of intermediates is still closely related to the final product morphology (**Figure 6C-6E**). The morphology-S-shapedness correlation across four lipid systems greatly helps rationalize the existence of non-standard vesicles.

Although numerous MD simulation studies have investigated the flake-to-vesicle transition^15,48–55,83^, S-shaped intermediates^51^ and non-standard products^49^ have rarely been reported. The most probable reason lies in two limitations of those studies. Previous studies typically limit the simulation system size to 100–3000 lipid molecules^15,17,48,49,52–55^, whereas our simulation system contains 6664 lipid molecules. The finite size effect of small simulation systems fails to capture long-wavelength undulations^84,85^ and can underestimate the coexistence of concave and convex regions, as well as the occurrence of non-standard products. Furthermore, previous studies also exhibit clear limitations in sampling, as the number of trajectories is limited to a few to a dozen^15,48,49,52–54^, especially for two studies that use larger systems containing 4800–5000 lipid molecules^50,83^. The inadequacy in sampling substantially reduces the probability of observing rare events, such as the formation of S-shaped intermediates and non-standard vesicles, let alone elucidating the relationship between them. Our work, however, generates hundreds of trajectories with a large simulation system, enabling a more systematic and comprehensive investigation.

Notably, further exploration of the flake-to-vesicle transition in larger systems is still warranted. Intermediates with topologies more complex than the S-shape may emerge, featuring multiple concave and convex regions. These intermediates could give rise to additional non-standard vesicular structures, such as FVCs composed of more than two daughter vesicles.

Based on our findings, we propose a generalized mechanism for the flake-to-vesicle transition that explicitly incorporates the formation of S-shaped intermediates and the emergence of non-standard vesicles such as SSVs, ELVs, and FVCs (**Figure 7**). During the initial BLF stage, the system samples a continuum of conformations with varying degrees of S-shapedness. Driven by edge tension, the membrane edge progressively contracts, which in turn modulates the S-shapedness of intermediates. Depending on the S-shapedness of these intermediates, four distinct product morphologies can arise: (1) intermediates with negligible or minor convex curvature evolve into RSVs; (2) those with a slightly enlarged convex region yield SSVs; (3) those with a larger convex region give rise to ELVs; and (4) those with a substantial convex proportion lead to FVCs. Notably, this new mechanism does not replace the classical one but generalizes it: for lipid systems that disfavor S-shaped intermediates, such as the pure DPSM and APM systems investigated in this work, the new mechanism largely reduces to the classical one. Given the conserved structural features across diverse vesicle-forming amphiphiles, this mechanism may not be limited to lipid vesicles, but may also be relevant to vesicles derived from surfactants^86–88^, block copolymers^89^, and hybrid assemblies^90^.

**Figure 7.**
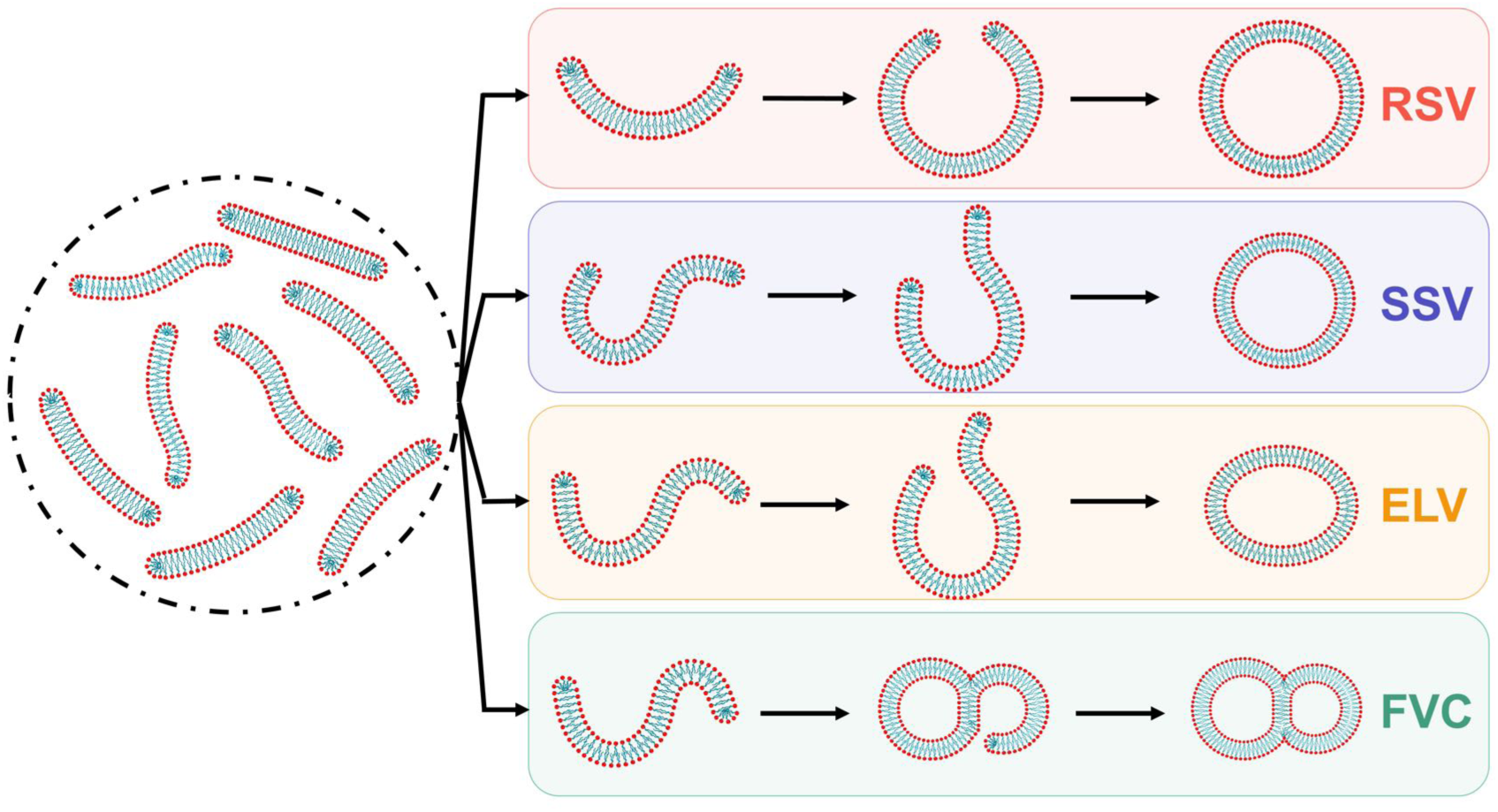
A generalized flake-to-vesicle mechanism that incorporates S-shaped intermediates and non-standard vesicles. Intermediates with varying degrees of S-shapedness lead to four distinct product morphologies, including RSV, SSV, ELV, and FVC.

Because product morphology depends on lipid composition and transbilayer distribution (**Table 1**) in our simulations, this dependence opens new opportunities for vesicle engineering. Experimental controllable parameters, such as lipid composition^91^, membrane protein content^92^, temperature^93,94^, solution type, and manufacturing procedure may be tuned to suppress or facilitate the formation of S-shaped intermediates and non-standard vesicles. Parameters that suppress the S-shaped intermediates and non-standard vesicles would yield a higher proportion of RSVs. This would contribute to minimizing product dispersity, which is crucial for reproducible drug loading, predictable in vivo performance, simplified quality control, and unambiguous interpretation of biophysical measurements^95,96^. Conversely, these parameters may be tuned to promote the formation of specific S-shaped intermediates and thereby selectively produce certain types of non-standard vesicles that may offer additional functionalities. Particularly, if FVCs are sufficiently stable, the separated compartments of the daughter vesicles may be exploited to load different types of cargoes independently, potentially providing a new multi-compartmentalized system for co-delivery of multiple drugs or synergistic agents^6,97^. These opportunities all stem from the generalized mechanism proposed here, in which the degree of S-shapedness of intermediates serves as a controllable determinant of the final product morphology.

## TOC

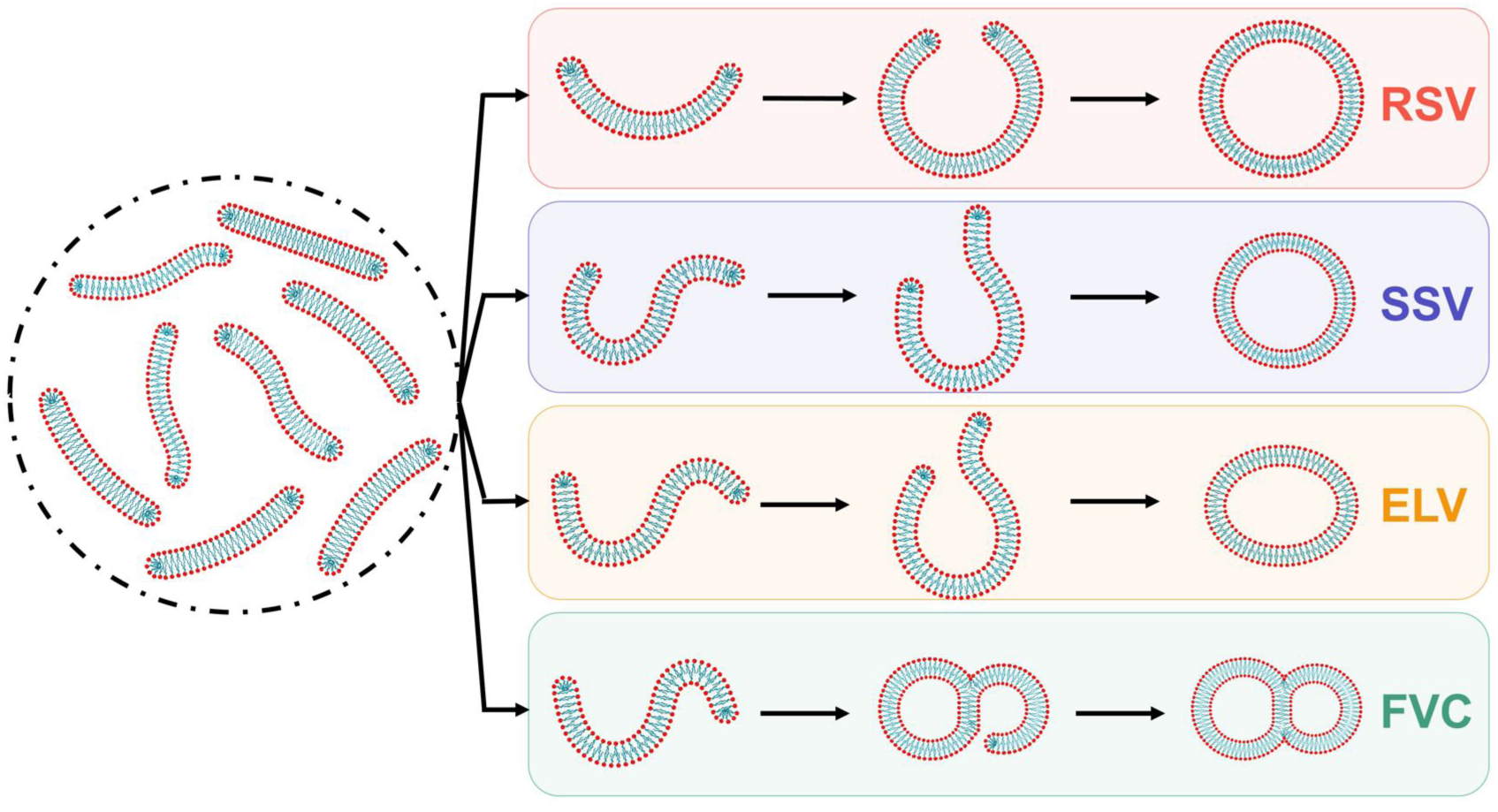

## Notes

### Competing Interest Statement

The authors have declared no competing interest.

### Funding

- National Key Research and Development Program of China: 2024YFA1208300
- National Natural Science Foundation of China: 22233002, 22573018

