## Supporting information for "Beyond the Classical Flake-to-Vesicle Mechanism: Non-standard Vesicles Arise from S-Shaped Intermediates"

**Preparation of lipid systems**

The pure 1,2-dioleoyl-sn-glycero-3-phosphoethanolamine (DOPE) system was first equilibrated as a planar bilayer in a rectangular simulation box with dimensions of 45.19 × 45.19 × 12.48 $nm^3$. Constructed using *insane*[1], the system comprised 6664 DOPE lipids and 142578 coarse-grained solvent beads, including 125548 W, 13950 WF, 1540 $Na^+$ and 1540 $Cl^-$ beads. After 500 ns of equilibrium, a 2-μs production run was performed under the semi-isotropic NPT ensemble. The trajectory was saved every 5 ns. 359 snapshots were extracted as starting structures for subsequent flake-to-vesicle transition simulations.

For each snapshot obtained above, the original solvent beads were removed to obtain a lipid flake structure. Each flake structure was then placed in a dodecahedron box with dimensions of 56.7 × 56.7 × 40.09 $nm^3$ and solvated by W and WF beads. $Na^+$ and $Cl^-$ beads substituted for a portion of the W/WF beads to achieve a final salt concentration of 0.15 M. The total number of beads in each bilayered flake system was maintained exactly at 976635. To minimize fluctuations in the box dimensions, each system was scaled to equilibrated dimensions and was subjected to energy minimization using the steepest descent algorithm for 20000 steps, followed by a 10-ns NVT equilibrium with position restraints applied to lipids using a force constant of 1000 $kJ \cdot mol^{-1} \cdot nm^{-2}$. An unrestrained production simulation of 200 ns was performed under the NPT ensemble with an isotropic pressure coupling. Totally 359 200-ns production trajectories were obtained for the DOPE system. One of the 200-ns trajectories initially ended in an incompletely formed ellipsoidal vesicle (ELV), and extending the trajectory to 1 μs allowed it to relax into a complete, stable ELV.

Three additional lipid systems, each containing 6664 lipids, were prepared following the same protocol. For the pure N-palmitoyl-D-erythro-sphingosyl-1-phosphocholine (DPSM) system, some production trajectories were extended to fully capture the complete vesiculation process. Each production trajectory of the asymmetric plasma membrane (APM) and the symmetric plasma membrane (SPM) systems lasted for 400 ns. The initial planar bilayer structure of the APM system composed of 63 distinct lipid species (encompassing 14 types of headgroups and 11 types of tails) was obtained from the Martini website (https://cgmartini.nl/docs/downloads/example-applications/lipid-membranes.html).

### Calculation of $A_p$ and SASA

The asphericity parameter ($A_p$) was employed to quantify the deviation of a three-dimensional object from perfect spherical symmetry. It is 1 for an extremely elongated or 'cigar' shaped object, 0.25 for a disk with infinitely small depth, and 0 for a perfect sphere, given by[2]:

$$A_p = \frac{(R_1^2 - R_2^2)^2 + (R_1^2 - R_3^2)^2 + (R_2^2 - R_3^2)^2}{2(R_1^2 + R_2^2 + R_3^2)^2}$$

where $R_i$ ($i$ = 1,2,3) are the eigenvalues of the gyration tensor $A$, which is given by:

$$A = \begin{bmatrix} S_{xx} & S_{xy} & S_{xz} \\ S_{yx} & S_{yy} & S_{yz} \\ S_{zx} & S_{zy} & S_{zz} \end{bmatrix}$$

where $S_{pp} = \frac{1}{n}\sum_i (x_i - x_c)^2$ and $S_{pq} = \frac{1}{n}\sum_i (p_i - p_c)(q_i - q_c)$ , $n$ is the total number of beads in the system, $p_i$ and $q_i$ ($p$, $q \in \{x, y, z\}$) denote the $p$- and $q$-coordinate of $i$th bead, respectively, and $p_c$ and $q_c$ are the corresponding coordinates of the center of mass of the system. Each bead has an equal weight in the calculation.

The solvent-accessible surface area (SASA) is defined as the surface traced by the center of a probe sphere representing a solvent molecule as it rolls over the molecular surface of interest. The SASA values for all lipids were computed using Gromacs built-in analysis tool (gmx sasa) with a probe radius of 0.264 nm. All van der Waals radii of the coarse-grained beads were set to the same value of 0.264 nm.

Automated analysis framework for curvature and boundary geometry of membranes **(ARCUS)**

To enable a direct comparison between MD snapshots and with the Helfrich elastic theory, we designed a novel method, termed the automated analysis framework for curvature and boundary geometry of membranes (ARCUS), to approximate a lipid bilayer as a 2-D midsurface (defined as the surface located midway between the two leaflets) and to calculate the mean curvature and the boundary edge length from the midsurface. The ARCUS approach comprises the following five steps:

**1. Selection of tail beads**

To construct the midsurface, the terminal beads of each lipid tail were selected as initial proxies, as they are generally located closest to the bilayer center. Two terminal-most beads from each lipid tail were chosen. In the DOPE lipid, these correspond to beads C3A and C4A from one tail, and C3B and C4B from the other. For the pure DSPM system, the corresponding beads are C3B and C4B from one tail and C2A and C3Afrom the other. For the plasma membrane systems, the bead selection is provided in a separate atom_selection_of_plasma_membrane.txt file (https://github.com/Zikang-Wang-FDU/Beyond-the-Classical-Flake-to-Vesicle-Mechanism).

**2. Generation of the bead cloud that defines the 2-D midsurface**

To obtain a zero-thickness midsurface, the selected tail beads from both leaflets were further compressed through an iterative displacement procedure. In each iteration, the new position of each tail bead was determined based on a bead collection consisting of the bead itself and its neighbors with a cutoff radius of $R_{\mathrm{MLS}}$ = 25 Å, a value found to perform well in this study. A covariance matrix was constructed from this bead collection, and its eigendecomposition yielded three eigenvectors. The eigenvector corresponding to the smallest eigenvalue was taken as the local surface normal. The tail bead was then projected onto the plane defined by this normal and the centroid of the bead collection, thereby updating its position. Beads with fewer than five neighbors were not displaced.

A maximum of 10 iterations were performed. However, the calculation was terminated early when the mean displacement of all beads fell below 0.05 Å. The final positions of the beads (*i.e.* the bead cloud) and their associated surface normals were subsequently used to define the 2-D midsurface.

#### 3. Construction of triangular surface mesh

The triangular surface mesh was constructed from the bead cloud using the ball-pivoting algorithm[3] (BPA) as implemented in the Open3D Library[4]. To accommodate variations in bead density across different regions of the cloud, three ball radii (5.0, 5.5, 6.0 Å) were sequentially applied during the BPA-based surface reconstruction. These radii were selected to maximize preservation of surface details (particularly the boundary edge) while minimizing the formation of isolated small triangles and mesh holes. The Open3D library was also used to orient all surface normals consistently toward the same side of the surface.

The resulting mesh was cleaned by merging duplicated vertices and removing unused beads. The largest connected component of the mesh was extracted to represent the lipid bilayer, while all smaller disconnected pieces were discarded.

#### 4. Calculation of the mean squared curvature of the entire mesh

The local mean curvature at each mesh vertex was computed using the algorithm implemented in the PyVista package[5]. The weight coefficient $w_i$ for vertex $i$ was defined as its associated surface area, calculated as the sum of the areas of all triangles containing vertex $i$, with ear triangle's area equally distributed among its three vertices:

$$w_i = \sum_{t \ni i} \frac{S_t}{3}$$

where $t$ denotes the $t$th triangle containing vertex $i$, and $S_t$ is the area of that triangle.

The mean squared curvature of the entire mesh was then defined as:

$$\langle H^2 \rangle = \sum_i H_i^2 \, w_i$$

To mitigate singularities in the surface mesh, the upper and lower 5% of $H_i$ values were trimmed to reduce the influence of extreme curvature outliers. This procedure yielded $\langle H^2 \rangle$ curves with no sharp spikes for the flake-to-vesicle trajectories (**Figure S2A**). This trimming step also reduced the S-shapedness index (SSI) from approximately 0.1 to approximately zero, while leaving the overall distribution of SSI almost unaffected.

5. Determination of the boundary edge of the 2-D surface

All mesh edges belonging to exactly one triangle were first identified, and a graph was constructed from these edges and their vertices. Since holes in the mesh can generate additional internal boundaries, all connected components of the graph were traversed, and the total perimeter of each component was computed. The component with the maximum perimeter was designated as the boundary edge of the 2-D midsurface.

**F1-score evaluation and threshold optimization**

To evaluate the correlation relationship of the S-shapedness of intermediates and the final product morphology, we used the SSI to quantify the S-shapedness of each intermediate, and assessed its ability to discriminate the final product morphology.

In a binary classification task (FVCs versus monomeric vesicles), a single threshold $\theta$ was applied to the SSI values (ranging from 0 to 1): structures with $0 \leq SSI < \theta$ were classified as monomeric vesicles, and those with $\theta \leq SSI \leq 1$ were classified as FVCs. The similarity between the predicted and ground-truth product morphologies are quantitatively evaluated using the macro-averaged F1-score, defined as:

$$F_{1,\mathrm{macro}} = \frac{1}{N}\sum_{i=1}^{N} F_{1,i}$$

where $N = 2$ and $F_{1,i}$ ($i \in$ {FVC, Monomeric}) are the pre-class binary F1-scores for FVCs and monomeric vesicles, respectively, in the binary classification task. No weighting factor is applied, ensuring that each morphology class contributes equally regardless sample size. For each binary F1-score, the positive class was defined as one morphology (either FVCs or monomeric vesicles), with the other serving as the negative class.

For a given positive class and the given threshold $\theta$, the following three counts are derived from the comparison between the predicted and ground-truth labels: true positives (TP) for which positive-class data correctly classified as positive, false positives (FP) for which negative-class data incorrectly classified as positive, and false negatives (FN) for which positive-class data incorrectly classified as negative.

Precision, the fraction of positive predictions that are correct, is defined by:

$$Precision = \frac{TP}{TP + FP}$$

and recall, the fraction of actual positive instances that are correctly identified, is defined by:

$$Recall = \frac{TP}{TP + FN}$$

and the binary F1-score was calculated as their harmonic mean:

$$F_1 = 2 \times \frac{Precision \times Recall}{Precision + Recall}$$

The F1-score was employed for evaluating classification performance because it provides a balanced measure of both precision and recall. Unlike accuracy, the F1-score robust to class imbalance, making it particularly suitable for datasets where one class may be underrepresented.

The threshold $\theta$ was then scanned across all candidate values, and the optimal value ($\theta^*$) was selected as the one that maximizes the macro-averaged F1-score.

To perform four-class classification of the final product morphologies using SSI as the sole metric, we defined three thresholds ($\theta_1$, $\theta_2$, $\theta_3$) over the SSI value with the following assignment rules: RSV for $0 \leq SSI \leq \theta_1$, SSV for $\theta_1 < SSI \leq \theta_2$, ELV for $\theta_2 < SSI \leq \theta_3$, and FVC for $\theta_3 < SSI \leq 1$. For each class $i \in$ {FVC, ELV, SSV, RSV}, a per-class binary F1-score ($F_{1,i}$) was computed by designating class $i$ as the positive class and the remaining three classes as the negative class. The three thresholds were jointly optimized by maximizing the macro-averaged F1-score across all four classes.

Two SSI variants were examined, corresponding to different stages of the flake-to-vesicle transition: (i) $SSI_{SFS}$, computed from five randomly extracted structures with SASA values of $7837 \pm 20\ \mathrm{nm}^2$ per trajectory in the pure DOPE system, and (ii) $SSI_{KPP}$, calculated from the structure immediately preceding the successful crossing of the key progress position in each trajectory.

High per-class and macro-averaged F1-scores at the optimized threshold indicates that the SSI possesses a strong discriminatory power between different product morphological classes.

**(A)**

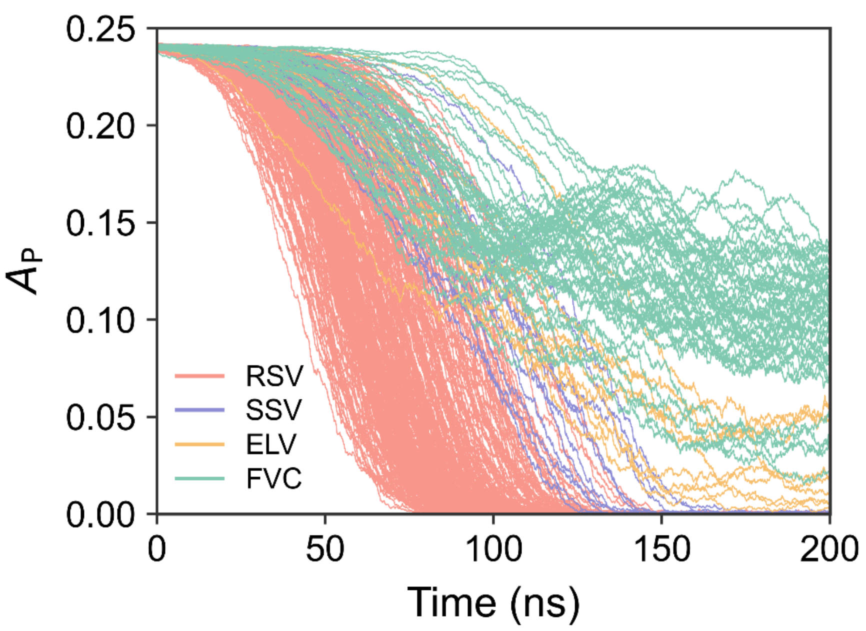


**(B)**

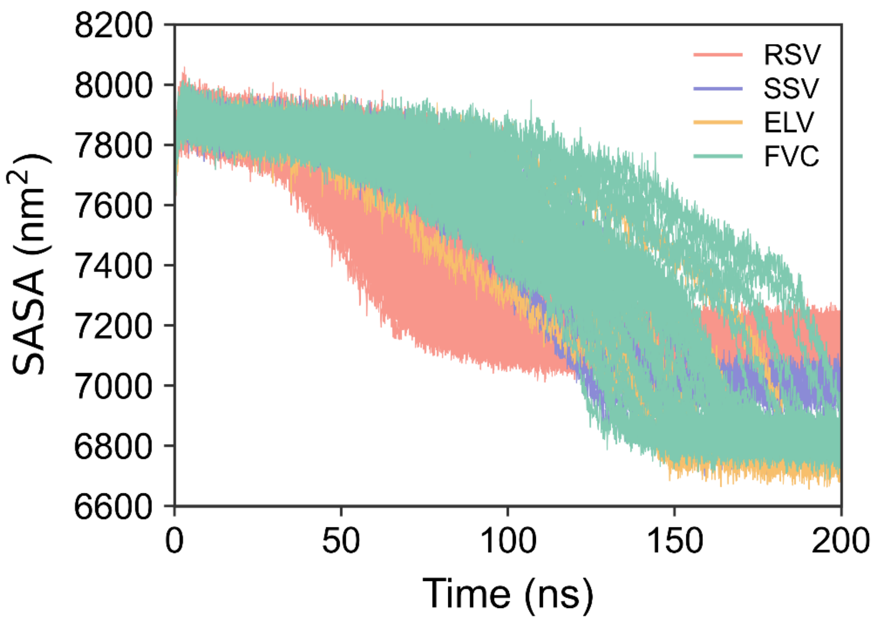


**Figure S1.** (A) Time evolution of the asphericity parameter ($A_P$) and (B) the solvent-accessible surface area (SASA) of all the lipids for each flake-to-vesicle trajectory of the pure DOPE system. In both panels, individual trajectories are colored according to the morphology of the corresponding final product.

**(A)**

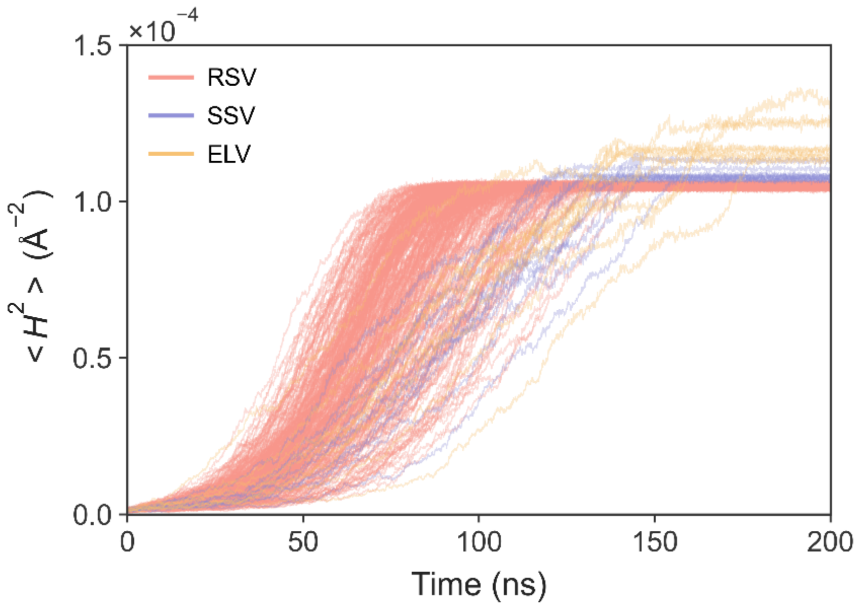


**(B)**

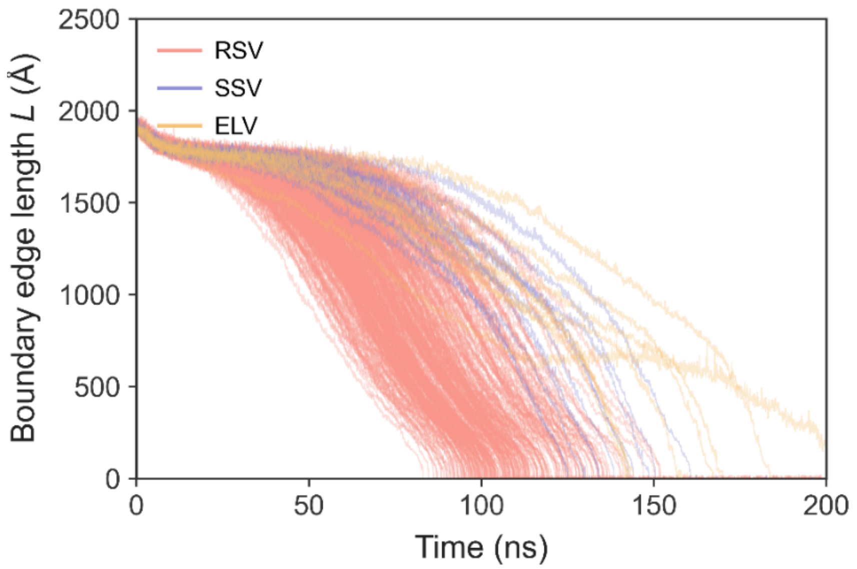


**Figure S2.** Time evolution of (A) the mean squared curvature ($<H^2>$) and (B) the boundary edge length of the bilayer, extracted from the flake-to-vesicle trajectories of the pure DOPE system that terminate in RSVs, SSVs and ELVs, respectively. Individual trajectories are colored according to the morphology of the corresponding final product. The singularities in the surface mesh were removed by trimming in the calculation of $<H^2>$. The three-way junction present in FVC-destined trajectories complicates the definition of a continuous 2-D midsurface, rendering the calculation of these quantities impractical within the current ARCUS approach. Therefore the data of FVC-destined trajectories are not shown.

(A)

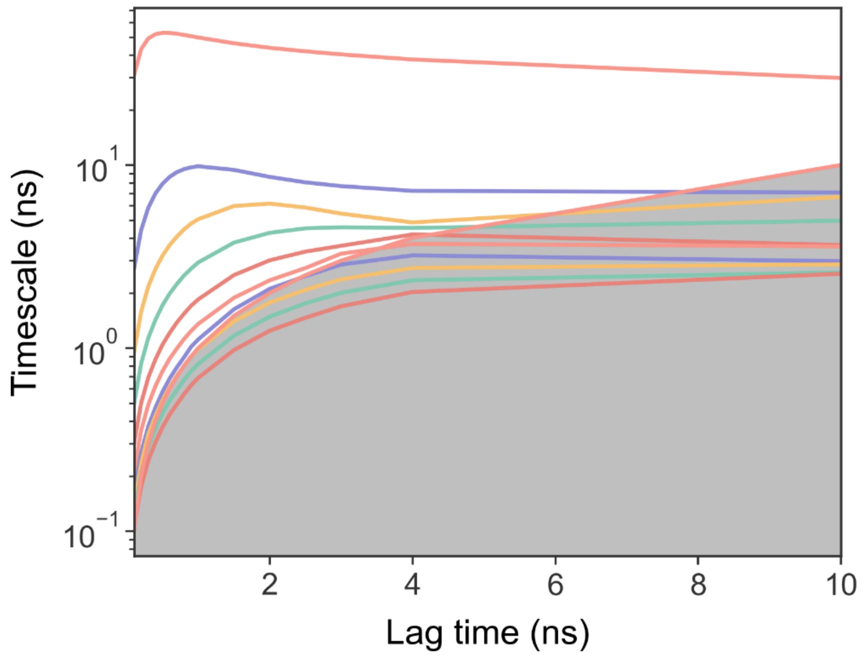

Timescale (ns)
$10^1$
$10^0$
$10^{-1}$
2
4
6
8
10
Lag time (ns)


(B)

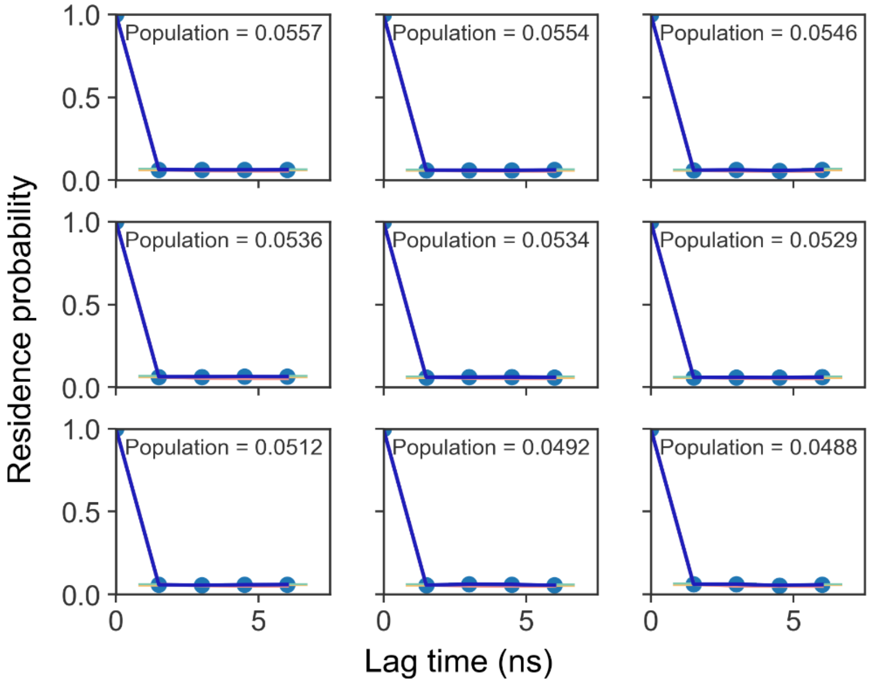

Residence probability
1.0
0.5
0.0
Population = 0.0557
Population = 0.0554
Population = 0.0546
Population = 0.0536
Population = 0.0534
Population = 0.0529
Population = 0.0512
Population = 0.0492
Population = 0.0488
0
5
Lag time (ns)


(C)

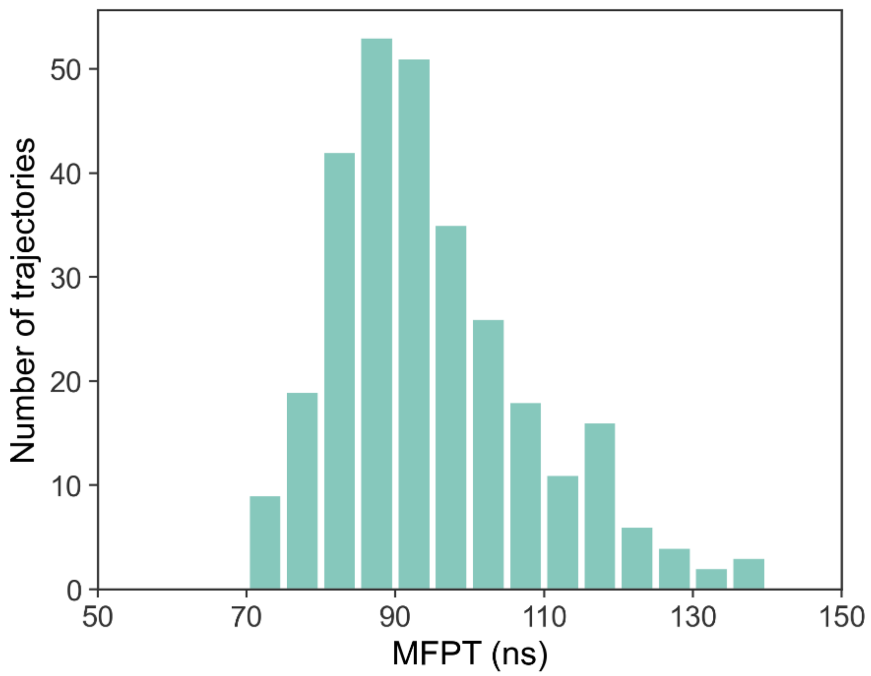

Number of trajectories
MFPT (ns)


(D)

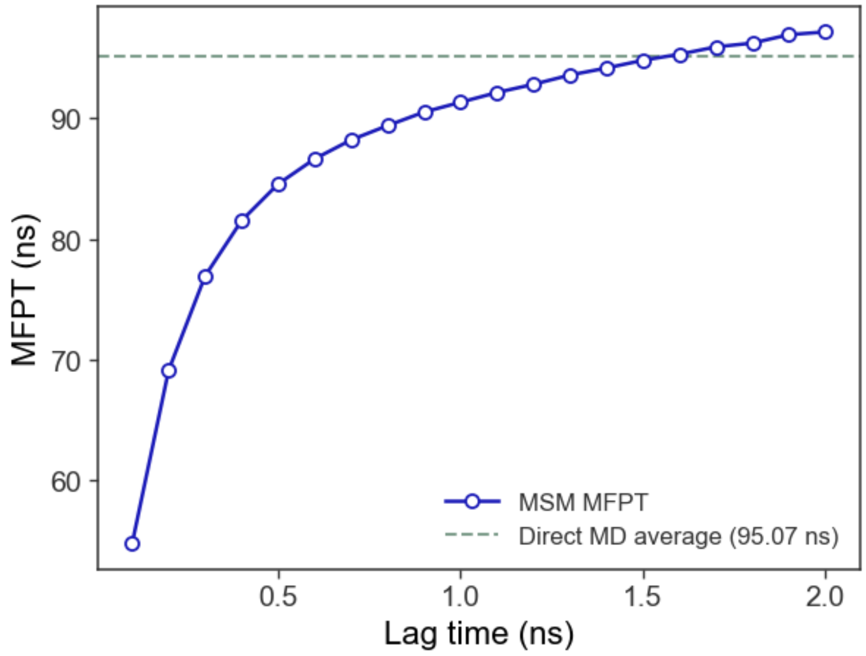

MFPT (ns)
Lag time (ns)
MSM MFPT
Direct MD average (95.07 ns)


‘

**(E)**

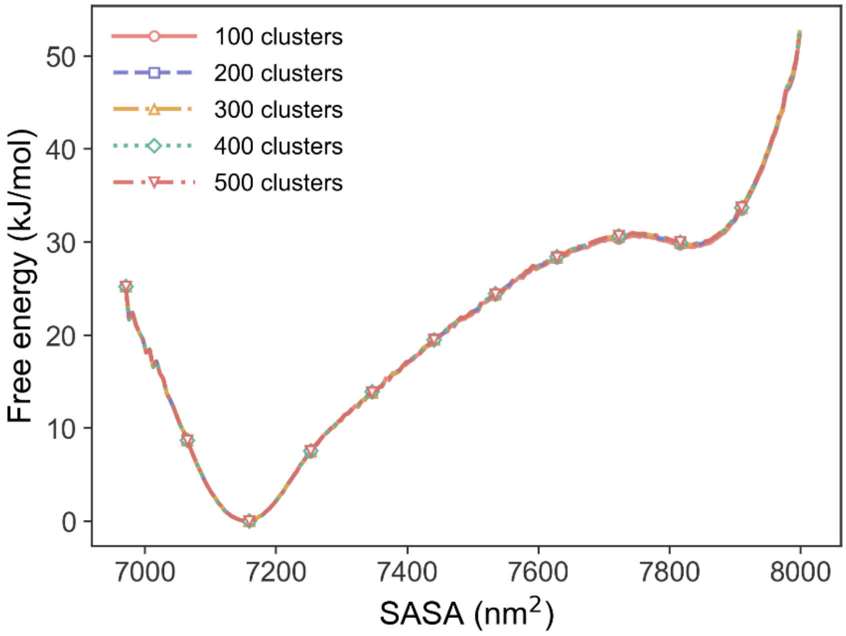


**Figure S3.** Validation of the 100-state Markov state model (MSM) constructed from the RSV-destined trajectories of the pure DOPE system. (A) Implied time scale test. (B) Chapman-Kolmogorov test (or residence test). (C) Distribution of first-passage time (FPT) extracted from the raw trajectories. FPT is defined as the time at which the SASA of lipids first drops below 7153 $nm^2$. The first 10 ns, discarded during MSM construction, were included in the FPT statistics. The mean first-passage time (MFPT) from the trajectories is 95.07 ns. (D) Dependence of the MFPT derived from MSM on the selection of lag time. The lag time of 1.5 ns was chosen as it yields a good agreement between the MSM-derived MFPT and the estimate directly from the MD trajectories. (E) One-dimensional free energy profiles derived from MSMs with varying numbers of microstates, showing that the profiles are largely insensitive to the number of microstates used.

(A)

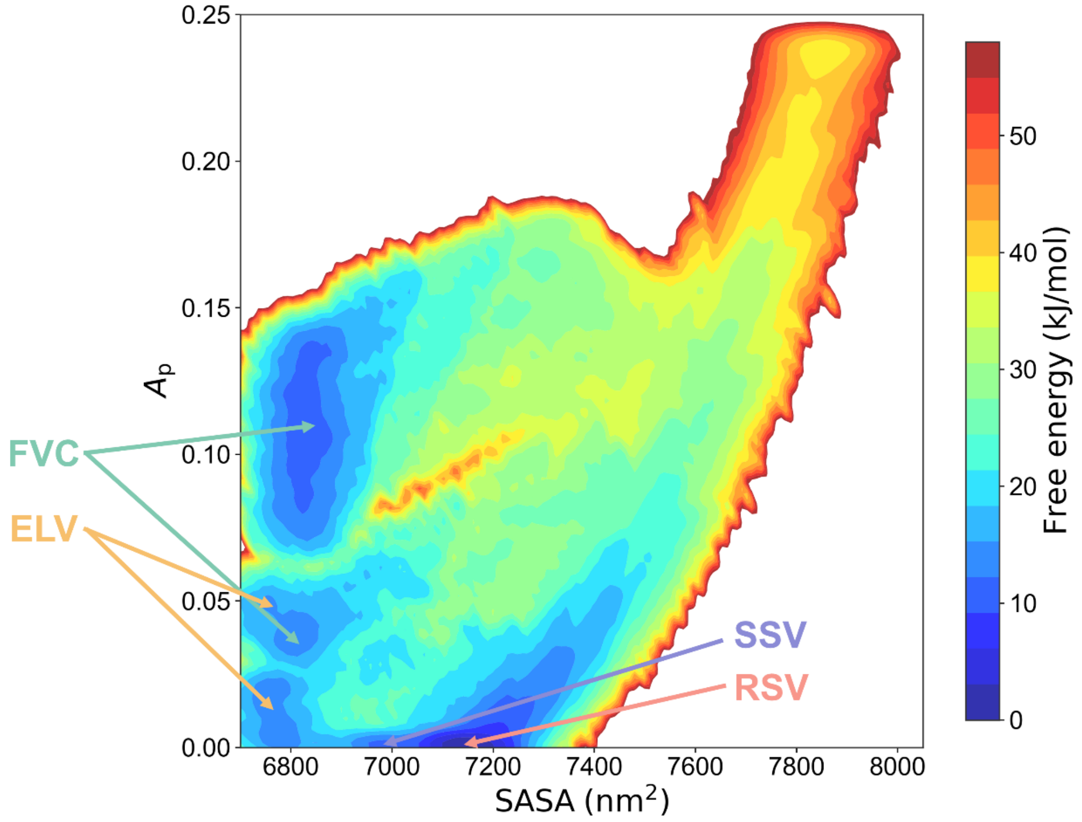

FVC
ELV
SSV
RSV
0.25
0.20
0.15
0.10
0.05
0.00
$A_p$
6800
7000
7200
7400
7600
7800
8000
SASA (nm$^2$)
Free energy (kJ/mol)
0
10
20
30
40
50


(B)

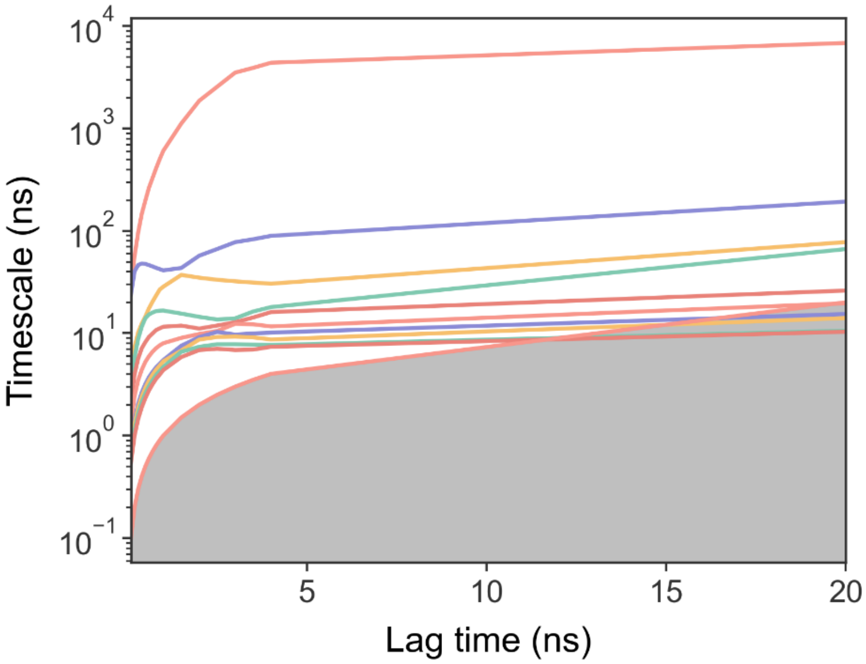

$10^4$
$10^3$
$10^2$
$10^1$
$10^0$
$10^{-1}$
Timescale (ns)
5
10
15
20
Lag time (ns)


(C)

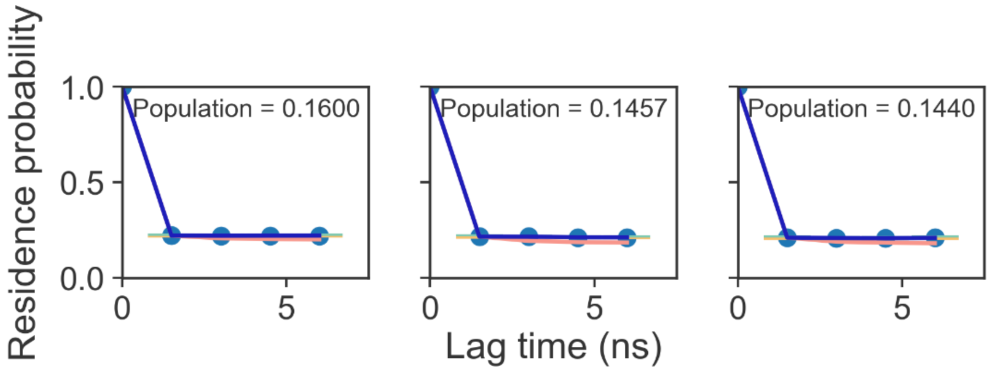


**Figure S4.** (A) Two-dimensional free energy landscape constructed using the lipid SASA and $A_p$ as collective variables, based on all 359 independent flake-to-vesicle trajectories of the pure DOPE system. Both $A_p$ and SASA were selected as features for clustering. Time-lagged independent component analysis (tICA) was first applied to these two variables using a lag time of 1.5 ns to recombine them, followed by K-means clustering of all the snapshots into 50 microstates. A lag time of 1.5 ns was chosen. The landscape reveals a global minimum corresponding to RSVs, a shallower local minimum associated with SSVs, two separated regions populated by ELVs, and two separated regions by FVCs. However, these two descriptors alone are insufficient to fully resolve the distinct formation pathways leading to the different final product morphologies. (B) Implied timescale test for the 50-state MSM. (C) Chapman-Kolmogorov test (or residence test).

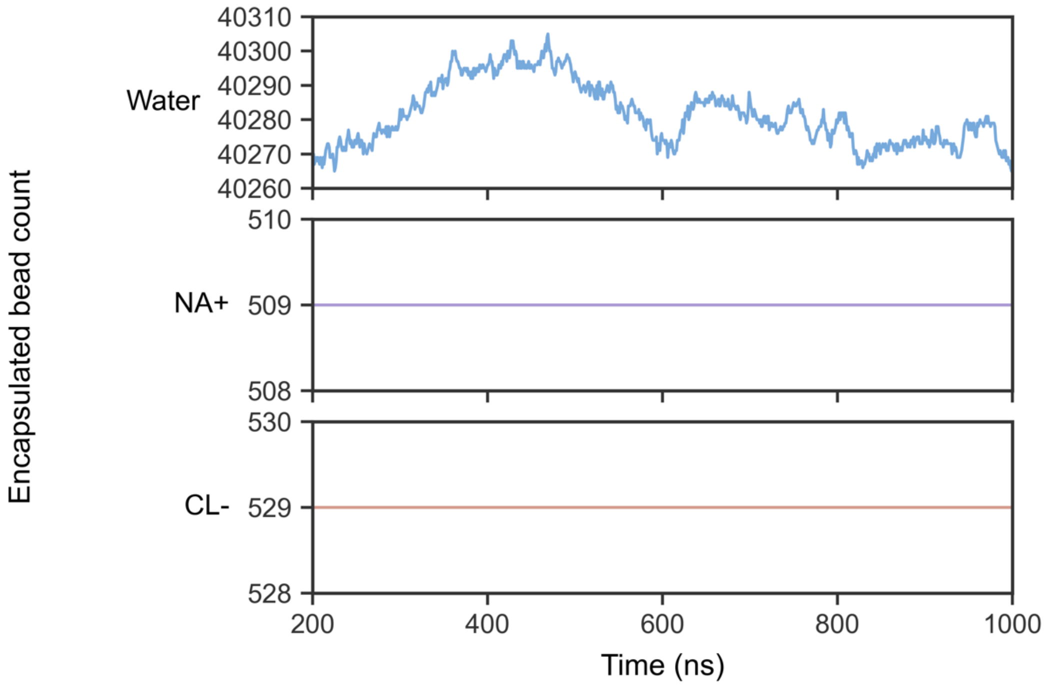


**Figure S5.** Solvent permeation across the lipid bilayer of a representative small-sized spherical vesicle (SSV) composed of pure DOPE, analyzed over the last 800 ns of a 1-μs trajectory. Time evolution of the number of (A) water beads, (B) $Na^{+}$ beads, and (C) $Cl^{-}$ beads enclosed within the vesicle cavity. Fluctuations in the number of water beads reflect their permeation through the lipid bilayer, whereas no permeation was observed for the ion beads.

(A)

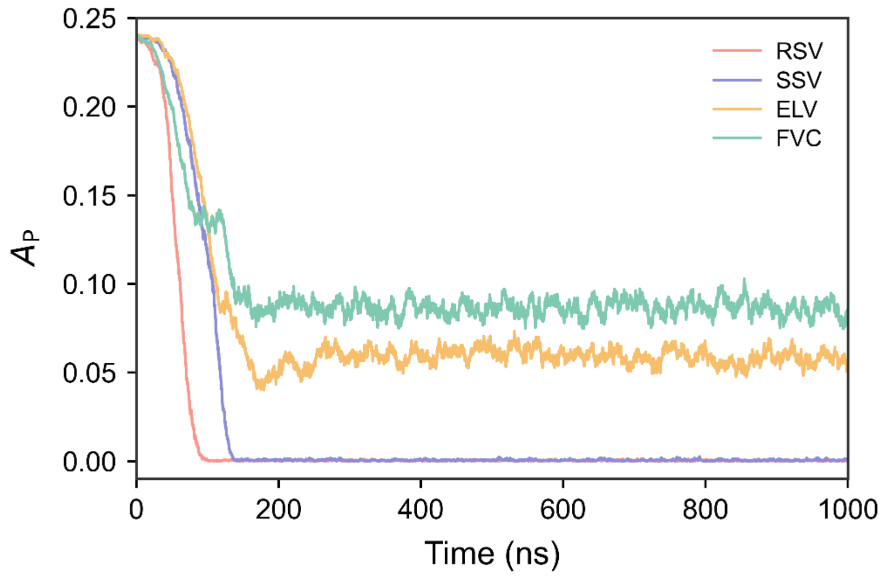

0.25
0.20
0.15
0.10
0.05
0.00
$A_P$
RSV
SSV
ELV
FVC
0
200
400
600
800
1000
Time (ns)


(B)

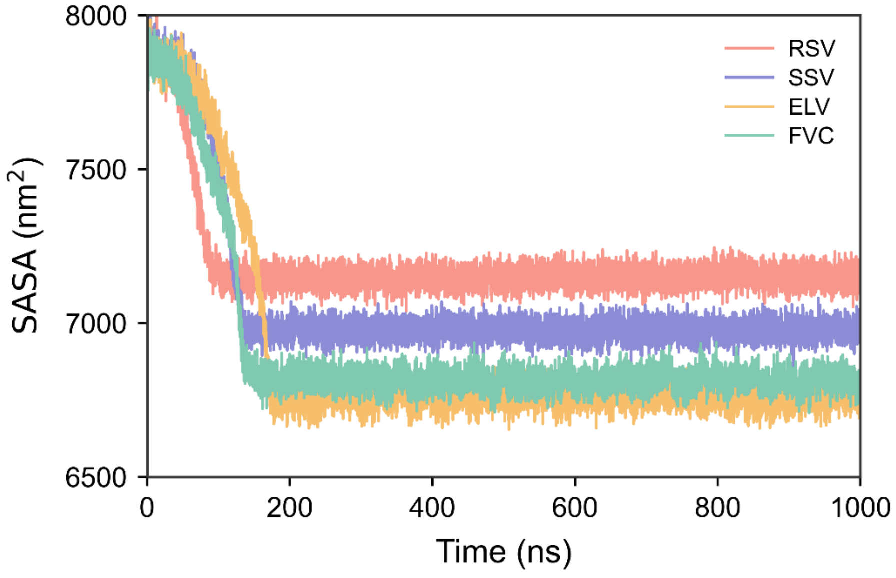

8000
7500
7000
6500
SASA (nm$^2$)
RSV
SSV
ELV
FVC
0
200
400
600
800
1000
Time (ns)

(C)

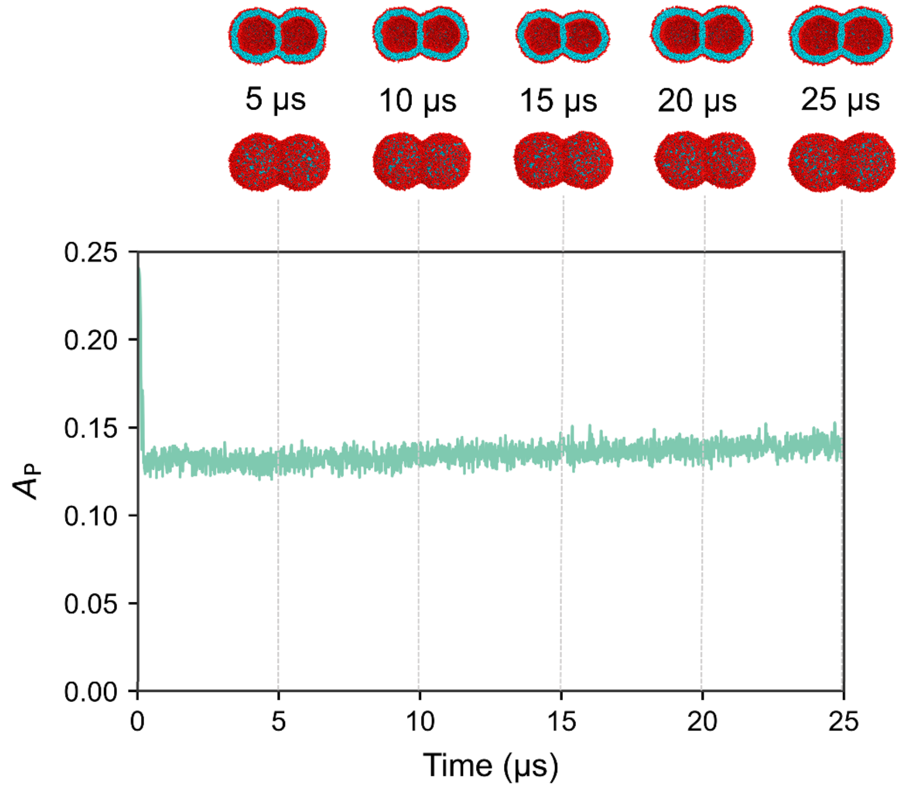


(D)

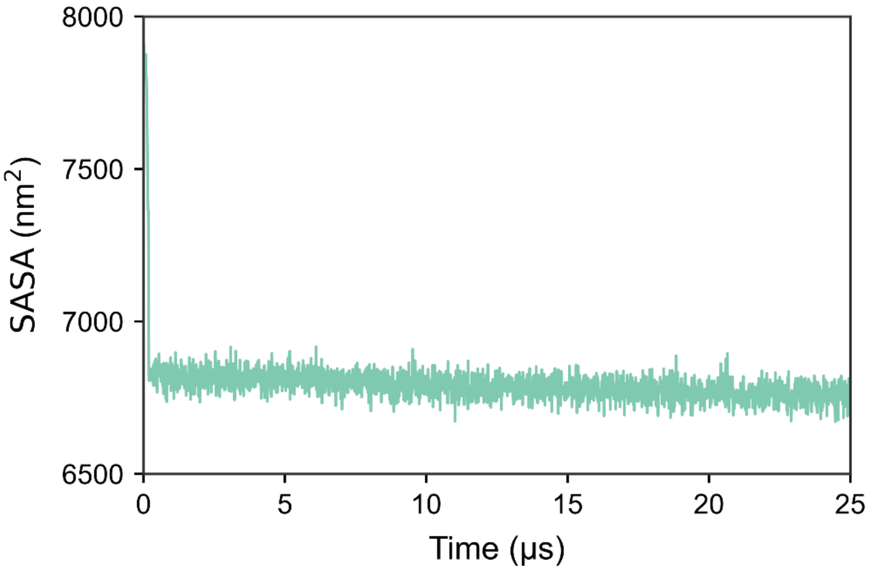


**Figure S6.** Extension of four representative 200-ns trajectories for the pure DOPE system, which ended in distinct final product morphologies. Time evolutions of (A) the asphericity parameter ($A_p$) and (B) the solvent-accessible surface area (SASA) of lipids during the extended simulation (to 1 μs). All product morphologies retained their structural integrity. Time evolutions of (C) $A_p$ and (D) lipid SASA for the FVC-destined trajectory when the simulation was extended to 25 μs. The actual simulation time of 25 μs corresponds to an effective 100 μs if a 4-fold timescale normalization is applied. The FVC slowly evolved toward a state with two daughter vesicles of comparable size.

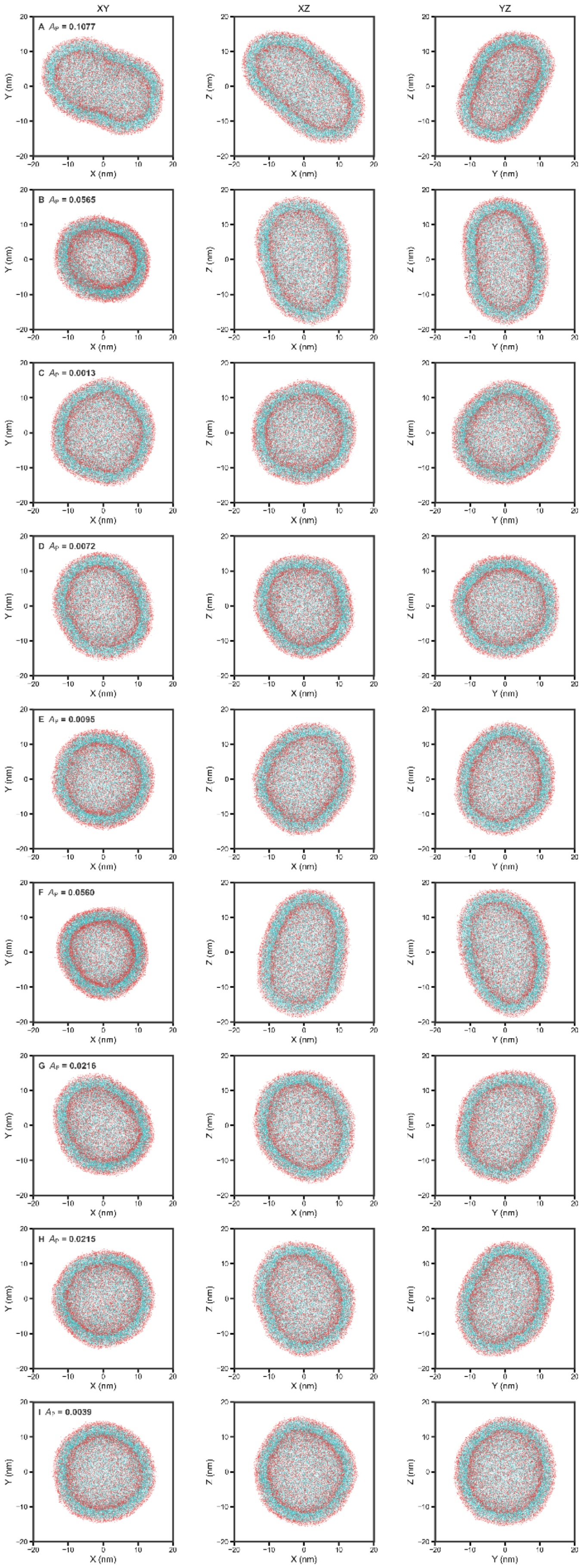

XY
XZ
YZ
A $A_P$ = 0.1077
B $A_P$ = 0.0565
C $A_P$ = 0.0013
D $A_P$ = 0.0072
E $A_P$ = 0.0095
F $A_P$ = 0.0560
G $A_P$ = 0.0216
H $A_P$ = 0.0215
I $A_P$ = 0.0039
X (nm)
Y (nm)
Z (nm)

**Figure S7.** Cross-section views of nine ELVs along three dimensions, obtained from the ELV-destined trajectories of the pure DOPE system. The $A_p$ values range from 0.0013 (quasi-spherical) to 0.1077 (highly elongated). All the ELVs are the final structures of the extended trajectories at 1 μs.

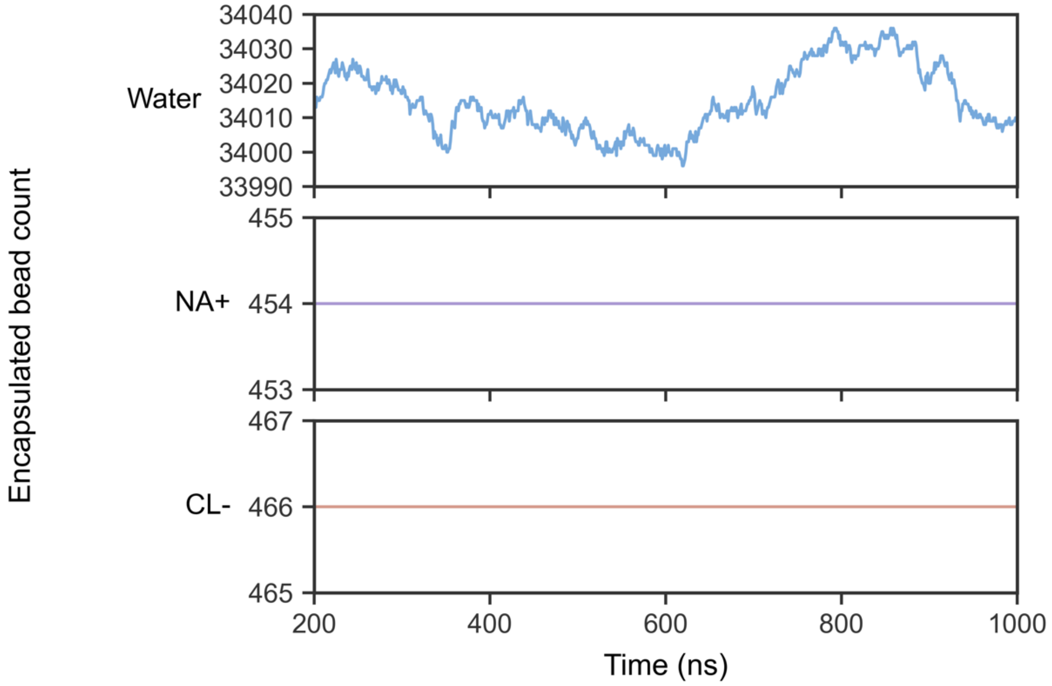


**Figure S8.** Solvent permeation across the lipid bilayer of a representative ellipsoidal vesicle (ELV) composed of pure DOPE, analyzed over the last 800 ns of a 1-μs trajectory. Time evolution of the number of (A) water beads, (B) $Na^+$ beads, and (C) $Cl^-$ beads enclosed within the vesicle cavity. Fluctuations in the number of water beads reflect their permeation through the lipid bilayer, whereas no permeation was observed for the ion beads.

(A)

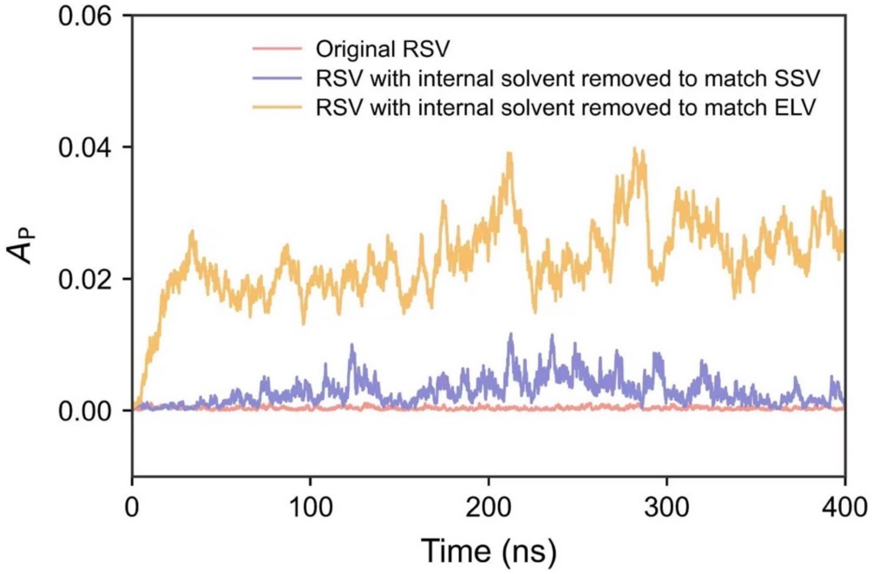


(B)

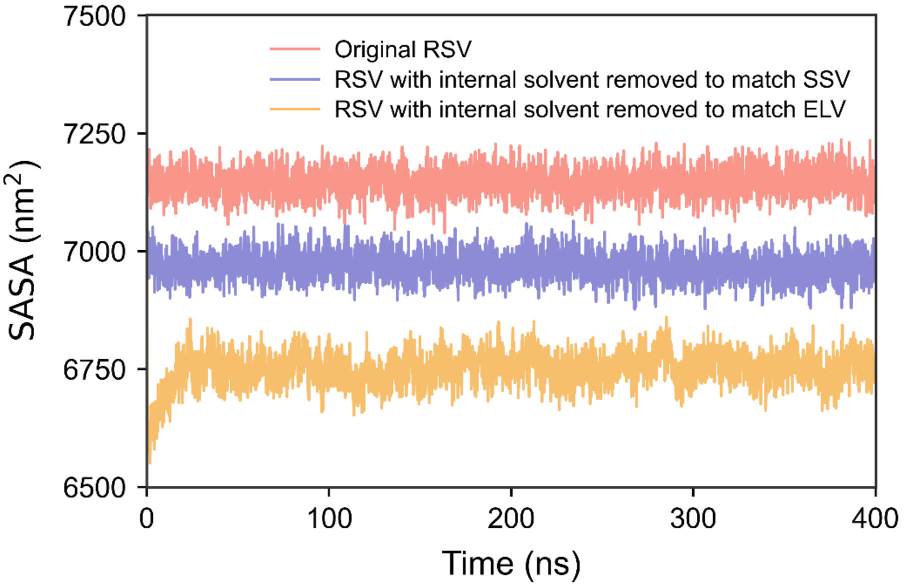


**Figure S9.** Transformation of an RSV composed of pure DOPE into an SSV or an ELV upon the removal of 2165 or 6689 internal solvent beads (water molecules or ions), respectively. The morphological changes are monitored by the time evolution of (A) $A_p$ and (B) the SASA. The trajectory with no solvent removed is also plotted for reference.

**(A)**

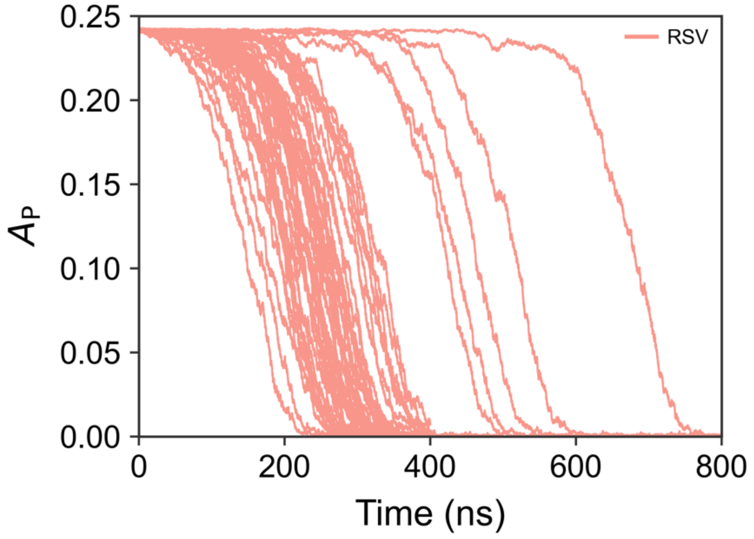


**(B)**

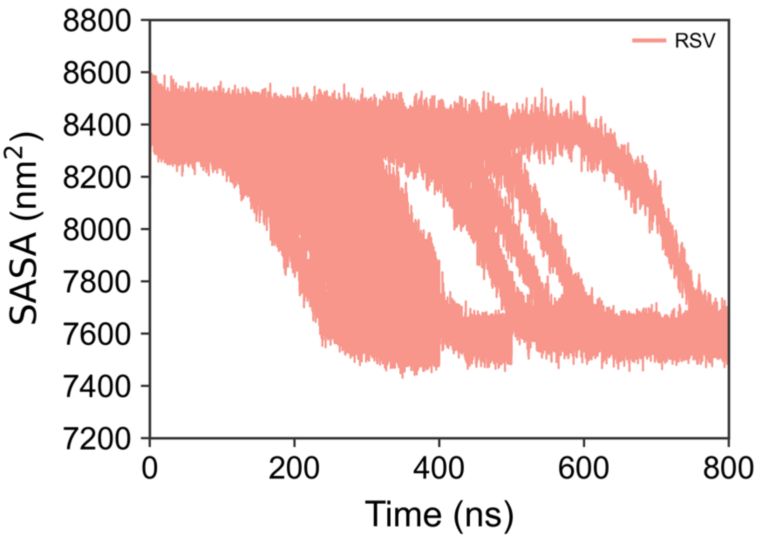


**Figure S10.** (A) Time evolution of the asphericity parameter ($A_p$) and (B) the solvent-accessible surface area (SASA) of all the lipids for each flake-to-vesicle trajectory of the pure DPSM system. All trajectories ended with an RSV.

(A)

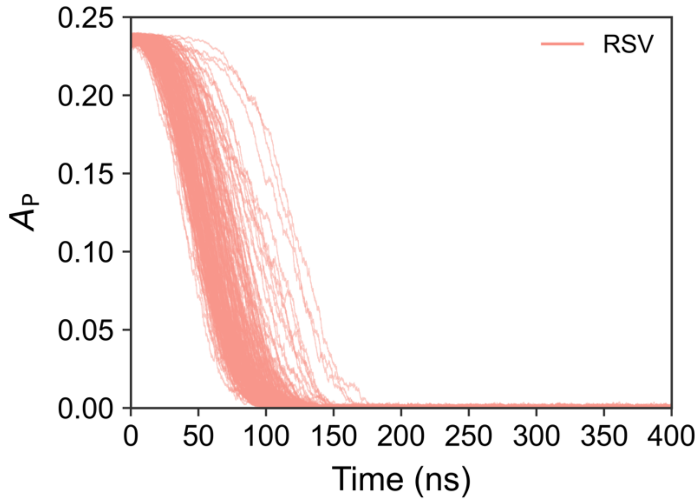


(B)

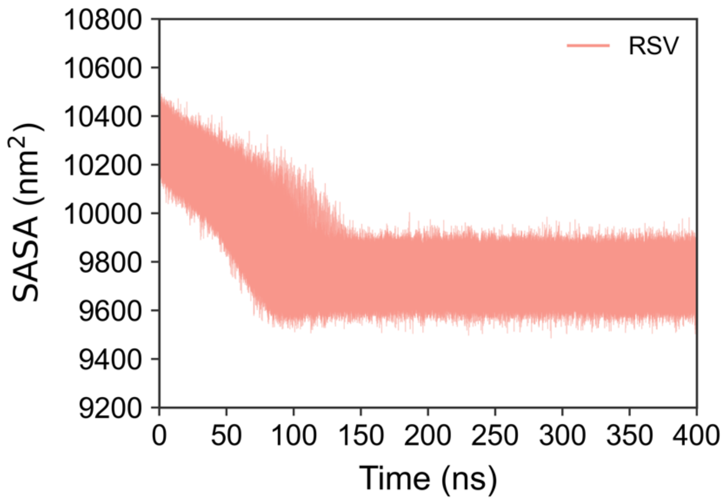


**Figure S11.** (A) Time evolution of the asphericity parameter ($A_p$) and (B) the solvent-accessible surface area (SASA) of all the lipids for each flake-to-vesicle trajectory of the asymmetric plasma membrane (APM) system. All trajectories ended with an RSV.

**(A)**

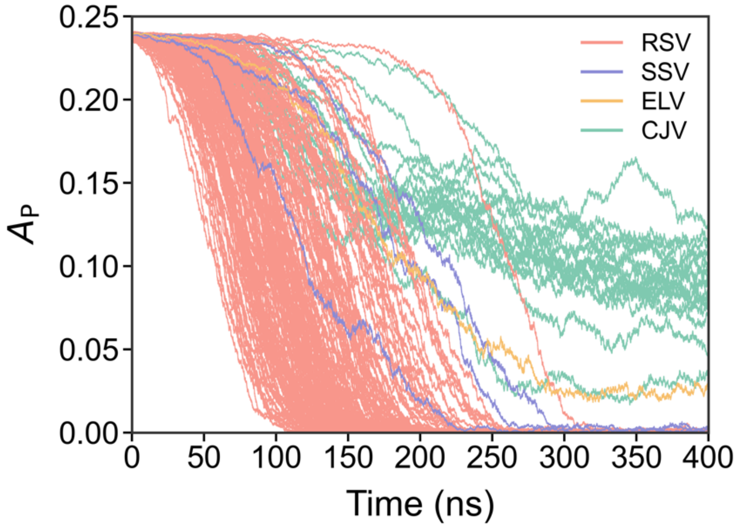

0.25
0.20
0.15
0.10
0.05
0.00
$A_P$
0
50
100
150
200
250
300
350
400
Time (ns)
RSV
SSV
ELV
CJV


**(B)**

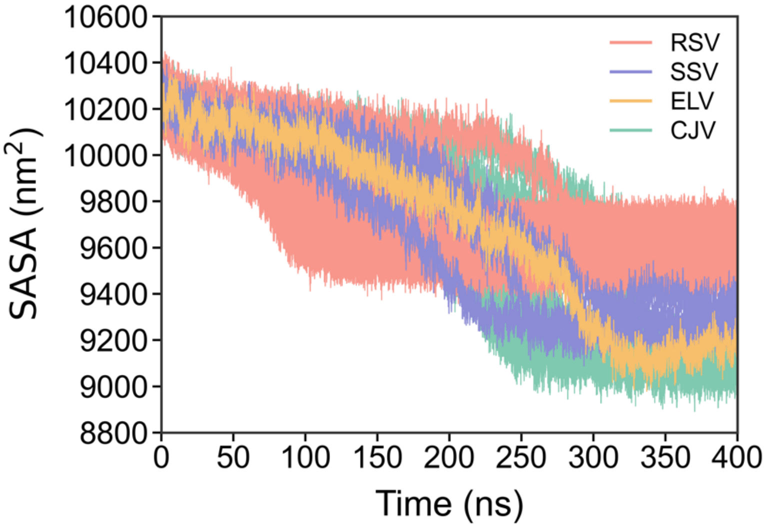

10600
10400
10200
10000
9800
9600
9400
9200
9000
8800
SASA (nm$^2$)
0
50
100
150
200
250
300
350
400
Time (ns)
RSV
SSV
ELV
CJV


**(C)**

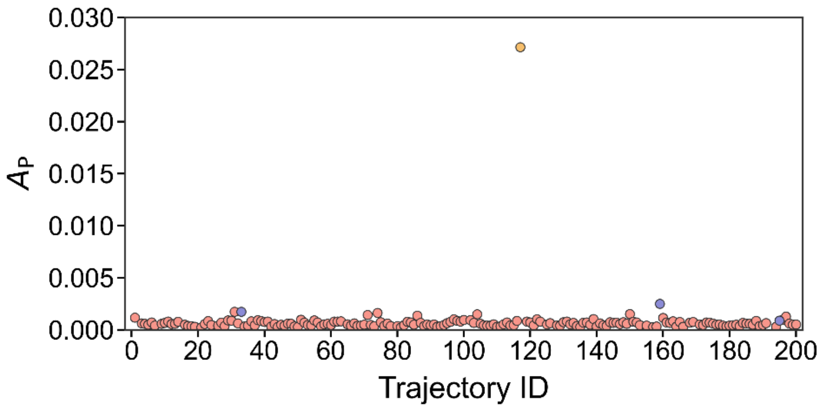

0.030
0.025
0.020
0.015
0.010
0.005
0.000
$A_P$
0
20
40
60
80
100
120
140
160
180
200
Trajectory ID


**(D)**

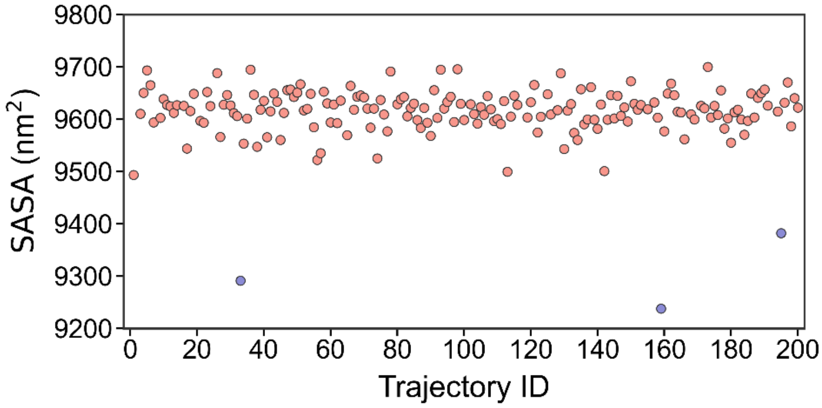


**Figure S12.** (A) Time evolution of the asphericity parameter ($A_\mathrm{p}$) and (B) the solvent-accessible surface area (SASA) of all the lipids for each flake-to-vesicle trajectory of the symmetric plasma membrane (SPM) system. In both panels, individual trajectories are colored according to the morphology of the corresponding final product. (C) Average $A_\mathrm{p}$ values of trajectories that ended with monomeric vesicles over the final 10 ns. The data point with an exceptionally high $A_\mathrm{p}$ (> 0.002, the orange point) corresponds to an ELV, whereas the remaining data points represent RSVs and SSVs. (D) Average SASA values of trajectories that ended with either RSVs and SSVs. The three data points (blue circles) with notably low SASA values correspond to SSVs, while the others correspond to RSVs.
